# B cell maturation restored ancestral germlines to control Omicron BA.2.86

**DOI:** 10.1101/2024.03.03.583187

**Authors:** Ida Paciello, Giulio Pierleoni, Elisa Pantano, Giada Antonelli, Piero Pileri, Giuseppe Maccari, Dario Cardamone, Giulia Realini, Federica Perrone, Martin Mayora Neto, Simone Pozzessere, Massimiliano Fabbiani, Francesca Panza, Ilaria Rancan, Mario Tumbarello, Francesca Montagnani, Duccio Medini, Piet Maes, Nigel Temperton, Etienne Simon-Loriere, Olivier Schwartz, Rino Rappuoli, Emanuele Andreano

**Affiliations:** Monoclonal Antibody Discovery (MAD) Lab, Fondazione Toscana Life Sciences, Siena, Italy; Department of Biotechnology, Chemistry and Pharmacy, University of Siena, Siena, Italy; Data Science for Health (DaScH) Lab, Fondazione Toscana Life Sciences, Siena, Italy; Viral Pseudotype Unit, Medway School of Pharmacy, Universities of Kent and Greenwich, Chatham Maritime, Kent, United Kingdom; Department of Cellular Therapies, Hematology and Laboratory Medicine, University Hospital of Siena, Siena, Italy; Department of Medical Sciences, Infectious and Tropical Diseases Unit, Siena University Hospital, Siena, Italy; Department of Medical Biotechnologies, University of Siena, Siena, Italy; KU Leuven, Rega Institute, Department of Microbiology, Immunology and Transplantation, Laboratory of Clinical and Epidemiological Virology, Leuven, Belgium; G5 Evolutionary Genomics of RNA Viruses, Institut Pasteur, Université Paris Cité, Paris, France; National Reference Center for Respiratory Viruses, Institut Pasteur, Paris, France; Virus and Immunity Unit, Department of Virology, Institut Pasteur, Paris, France; Vaccine Research Institute, Creteil, France; Fondazione Biotecnopolo di Siena, Siena, Italy

**Author notes:** Corresponding author: Emanuele Andreano.

## Abstract

The unceasing interplay between SARS-CoV-2 and the human immune system has led to a continuous maturation of the virus and B cell response providing an opportunity to track their evolution in real time. We longitudinally analyzed the functional activity of almost 1,000 neutralizing human monoclonal antibodies (nAbs) isolated from vaccinated people, and from individuals with hybrid and super hybrid immunity (SH), developed after three mRNA vaccine doses and two breakthrough infections. The most potent neutralization and Fc functions against highly mutated variants, including BA.2.86, were found in the SH cohort. Despite different priming, epitope mapping revealed a convergent maturation of the functional antibody response. Neutralization was mainly driven by Class 1/2 nAbs while Fc functions were induced by Class 3/4 antibodies. Remarkably, broad neutralization was mediated by restored IGHV3-53/3-66 B cell germlines which, after heterogenous exposure to SARS-CoV-2 S proteins, increased their level of somatic hypermutations. Our study shows the resilience of the human immune system which restored previously expanded germlines and activated naïve B cells to broaden the antibody repertoire of antibodies to control future SARS-CoV-2 variants.

## INTRODUCTION

The severe acute respiratory syndrome coronavirus 2 (SARS-CoV-2) has shown an incredible ability to evolve and evade the antibody response elicited after infection and vaccination. Since the beginning of the coronavirus disease 2019 (COVID-19) pandemic, over seventy variants under monitoring, of interest and of concern have emerged highlighting the great plasticity of this virus and the ability to adapt to the human immune response over time^1^. Indeed, the initial antibody response induced by SARS-CoV-2 infection was dominated by IGHV3-53/3-66 encoded antibodies which were able to potently neutralize the original virus isolated in Wuhan, China, in their germline-like state^2–8^. These antibodies are known to target the spike (S) protein receptor binding domain (RBD), which is overall the main target for SARS-CoV-2 neutralizing antibodies (nAbs)^8–10^. Given the high immune pressure generated by these germlines, the first variants that rapidly emerged and spread worldwide, like the B.1.1.7, B.1.351 and B.1.1.248, introduced the E484K and K417N mutations that showed to evade almost 60% of antibodies encoded by IGHV3-53/3-66 germlines in both infection and mRNA vacciantion^2,7^. The subsequent waves of the pandemic were driven by SARS-CoV-2 variants with increased number of mutations in the S protein. Specifically, the appearance of the first Omicron variants (BA.1 and BA.2) at the end of 2021 marked a new chapter in the COVID-19 pandemic^11,12^. The BA.1 and BA.2 variants harbored 37 and 31 mutations in the S protein showing unprecedented immune evasion levels reducing drastically the neutralizing efficacy of sera from infected and vaccinated people and evading almost 90% of IGHV3-53/3-66 gene derived nAbs^13–16^. From 2022 onward, several Omicron variants appeared worldwide with increased number of mutations and immune evasion levels starting from the BA.5 to BA.2.86, the most mutated variant ever observed carrying almost 60 mutations in the S protein^17–24^. These variants kept spreading and infecting people worldwide generating a more mature and broadly reactive antibody response to SARS-CoV-2. To understand how the immunological response adapted and matured to SARS-CoV-2 variants over time, we longitudinally analyzed at single-cell level almost 1,000 nAbs isolated from four different cohorts: seronegative donors that received two mRNA vaccine doses (SN2); the same donors in SN2 subsequently re-enrolled when received a third booster dose (SN3); seropositive subjects with hybrid immunity, i.e. one infection and two mRNA vaccine doses (SP2); seropositive donors with super hybrid immunity (SH), i.e. at least three mRNA vaccine doses and two breakthrough infection. Part of donors in this cohort were re-enrolled from the SP2 group. Our study revealed that SH had the strongest antibody response against all tested variants. The cross-protective response was dominated by highly mutated IGHV3-53/3-66 gene-derived nAbs that, after additional infection and vaccination, restored their neutralization and Fc function activities against all variants including the highly mutated BA.2.86. Noteworthy, a strong convergence in the Class of RBD-targeting nAbs and germline expanded in SN3 and SH was observed with nAbs isolated in this latter cohort showing higher levels of cross-protection. Anyway, despite high breadth to SARS-CoV-2 variants, we observed that cross-reactivity was not extended to other alpha or beta human coronaviruses in the SH cohort. In addition, SH activated new germlines to broaden the antibody repertoire with new B cell germlines which could be rapidly deployed with the emergence of future SARS-CoV-2 variants. Taken together, this work dissects the functional antibody response currently developed in most people and identifies the immunological and genetic features behind cross-protection to highly mutated SARS-CoV-2 variants.

## RESULTS

### Frequency of B cells and neutralizing antibodies in super hybrid immunity

In this study we evaluated the B cell maturation and neutralizing antibody response of donors with super hybrid immunity (SH), i.e., at least three mRNA vaccine doses and two breakthrough infections. Six donors were analyzed in this work, two of which (VAC-004 and VAC-009) were re-enrolled from our previous study where we evaluated their B cell and antibody response following two mRNA vaccine doses and one breakthrough infection (hybrid immunity)^7^. Blood collection for SH donors occurred at an average of 99 days after the last vaccination dose or SARS-CoV-2 breakthrough infection. SH subject details are summarized in **Supplementary Table 1.** To evaluate the breadth of reactivity of CD19^+^CD27^+^IgD^-^IgM^-^ class-switched memory B cell (MBCs) towards different betacoronaviruses, we stained the cells with the Spike (S) protein of both SARS-CoV-1 and SARS-CoV-2 Wuhan and analyzed the frequencies of single and double positive cells (**Supplementary Fig. 1a**). We selected the SARS-CoV-2 Wuhan S protein as staining bait as all donors were exposed to this antigen. As expected, SH donors showed the highest frequency towards the S protein of SARS-CoV-2 with an average of 0.32% of positive cells, followed by B cells reactive to the SARS-CoV-1 S protein (0.13%) (**Supplementary Table 2**). Double positive MBCs showed the lowest frequency averaging 0.03% of reactive cells. To evaluate at single level the neutralizing antibody response of SH donors, SARS-CoV-1 and SARS-CoV-2 Wuhan S protein single and double positive MBCs were single cell sorted and incubated for two weeks to naturally release human monoclonal antibodies (mAbs) into the supernatant. A total of 4,505 S protein^+^ MBCs were sorted and directly tested in neutralization against the live SARS-CoV-2 Wuhan virus and SARS-CoV-1 pseudovirus. Overall, 9.4 to 16.3% of antibodies from the six SH donors showed capacity to neutralize at least one virus, resulting in a panel of 545 (12.1%) neutralizing human monoclonal antibodies (nAbs) (**Supplementary Fig. 1b, bottom panel; Supplementary Table 2**). The fraction of neutralizing antibodies is comparable to what was previously observed for the SN3 (14.4%) and SP2 (14.8%) cohorts^7,25^. Of these, 507 (93.0%), 27 (5.0%) and 11 (2.0%) were able to neutralize live SARS-CoV-2 Wuhan virus, SARS-CoV-1 pseudovirus or both betacoronaviruses respectively (**Supplementary Fig. 1b, top panel; Supplementary Table 2**).

### Antibody neutralization to SARS-CoV-2 Omicron variants

To better characterize identified nAbs, we tried to express all 545 as immunoglobulin G 1 (IgG1) and recovered 419 of them. Of these, 410 (97.8%) neutralized SARS-CoV-2 Wuhan, 7 (1.7%) were cross-neutralizing, and 2 (0.5%) neutralized only SARS-CoV-1. To evaluate their potency and breadth to SARS-CoV-2 and its variants, 417 nAbs (SARS-CoV-2 only and cross-neutralizing) isolated from SH individuals were tested by cytopathic effect-based microneutralization assay (CPE-MN) against SARS-CoV-2 Wuhan and Omicron BA.5, BA.2.75, BF.7, BQ.1.1, XBB.1.5, EG.5.1.1 and BA.2.86 variants. The data obtained from SH donors were compared with donors seronegative to SARS-CoV-2 infection but vaccinated with two (SN2; n=5) or three (SN3; n=4) mRNA vaccine doses, and with seropositive subjects with hybrid immunity (two mRNA vaccine doses and one breakthrough infection; SP2; n=5) (**Fig. 1a-d; Supplementary Fig. 1c-d; Supplementary Fig. 2a-d**). From SN2, SN3 and SP2, we previously isolated 52, 206 and 224 nAbs respectively^7,25^. Neutralization potency was expressed as 100% inhibitory concentration (IC_100_) and the evaluation of all antibodies in each cohort as geometric mean IC_100_ (GM-IC_100_). SH donors had an overall higher percentage of nAbs neutralizing all SARS-CoV-2 Omicron variants (**Fig. 1a-d; Supplementary Table 3**). Only one nAb (1.9%) was able to neutralize BF.7 and BQ.1.1 in the SN2 while none of the antibodies in this cohort showed neutralization activity against the other Omicron variants tested (**Fig. 1a; Supplementary Fig. 2a**). In contrast, higher levels of cross-protection were observed in SN3, SP2 and SH. The frequencies of nAbs from SN3 donors neutralizing Omicron BA.5, BA.2.75, BF.7, BQ.1.1, XBB.1.5, EG.5.1.1 and BA.2.86 variants were 12.1 (n=23), 17.0 (n=32), 10.2 (n=21), 5.8 (n=15), 7.3 (n=14), 3.4 (n=7) and 2.9% (n=6), while these variants were neutralized by 13.4 (n=30), 3.6 (n=8), 9.8 (n=22) and 5.8% (n=13) of SP2 antibodies (**Fig. 1b-c; Supplementary Fig. 2b-c**). None of the nAbs in the SP2 cohort were able to neutralize XBB.1.5, EG.5.1.1 and BA.2.86 variants. Finally, the frequencies of nAbs from SH donors neutralizing BA.5, BA.2.75, BF.7, BQ.1.1, XBB.1.5, EG.5.1.1 and BA.2.86 variants were 62.1 (n=259), 22.5 (n=94), 42.2 (n=175), 34.3 (n=143), 18.7 (n=76), 6.5 (n=27) and 14.4% (n=60) respectively (**Fig. 1d; Supplementary Fig. 2d**). The four cohorts showed similar neutralization potencies against SARS-CoV-2 Wuhan, nAbs isolated from SH donors had the highest neutralization potencies against all Omicron variants tested and it was the only cohort with nAbs showing a potency below 10 ng ml^-1^ (**Fig. 1a-d; Supplementary Fig. 1c-d; Supplementary Table 3-4**). To understand the S protein domains targeted by antibodies isolated from SH donors, nAbs were tested for binding against the receptor binding domain (RBD), N-terminal domain (NTD) and the S2 domain of the original Wuhan SARS-CoV-2 S protein. In all SH subjects, nAbs targeted mainly the RBD (n=317; 75.3%) followed by NTD (n=81; 19.2%) and S protein (n=23; 5.5%) (**Supplementary Fig. 2e; Supplementary Table 4**). No S2 domain binding nAbs were identified. This distribution is in line to what was previously observed in donors with three mRNA vaccine doses and with hybrid immunity^25^.

**Fig. 1.**
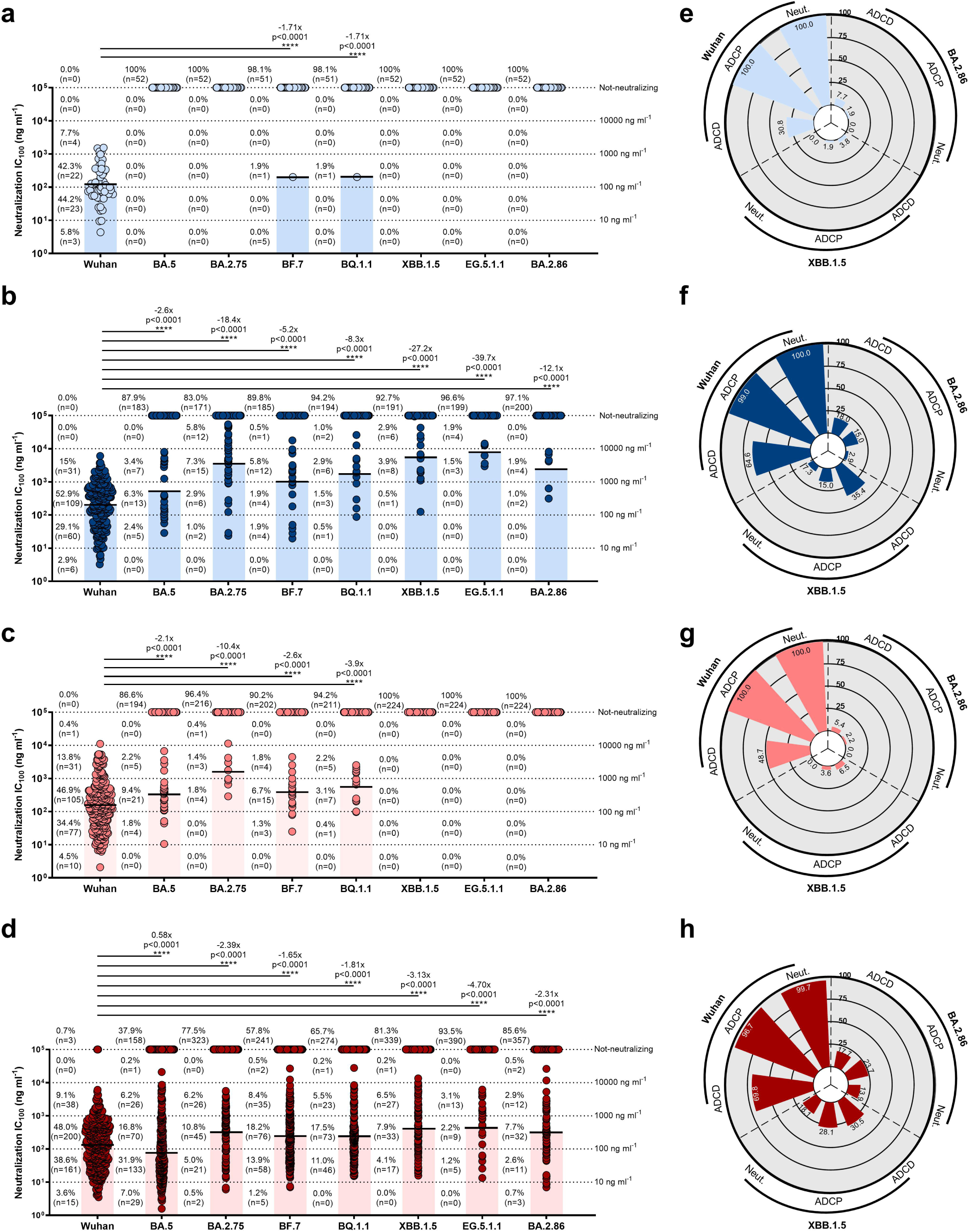
nAbs potency and breadth of neutralization and Fc functions against SARS-CoV-2 Omicron variants. **a-d,** Scatter dot charts show the neutralization potency, reported as IC_100_ (ng ml^-1^), of nAbs tested against the original Wuhan SARS-CoV-2 virus, and the Omicron BA.5, BA.2.75, BF.7, BQ.1.1, XBB.1.5, EG.5.1.1 and BA.2.86 lineages for SN2 (**a**), SN3 (**b**), SP2 (**c**) and SH (**d**). The number, percentage, GM-IC_100_ (black lines and colored bars), fold-change and statistical significance of nAbs are denoted on each graph. Reported fold-change and statistical significance are in comparison with the Wuhan virus. Technical duplicates were performed for each experiment. **e-h**, Doughnut charts show the frequency of nAbs retaining neutralization (Neut.), ADCP and ADCD against the SARS-CoV-2 Wuhan virus and the XBB.1.5 and BA.2.86 variants for SN2 (**e**), SN3 (**f**), SP2 (**g**) and SH (**h**). A nonparametric Mann–Whitney t test was used to evaluate statistical significances between groups. Two-tailed p-value significances are shown as *p < 0.05, **p < 0.01, and ***p < 0.001.

### Evaluation of Fc effector functions to XBB.1.5 and BA.2.86

Next, we evaluated the antibody-dependent phagocytic activity (ADCP) and antibody-dependent complement deposition (ADCD), of all identified nAbs in SH against the ancestral Wuhan virus, the XBB.1.5 variant that dominated from February to May 2023, and the highly mutated SARS-CoV-2 variant BA.2.86, ancestral germline of the currently predominant JN.1 variant (**Fig. 1e-h**). The Fc functions were evaluated for nAbs that retained the binding to the XBB.1.5 and BA.2.86 S proteins (**Supplementary Fig. 3a**). Binding nAbs were 7.7 (n=4), 43.7 (n=90), 11.6 (n=26) and 39.8% (n=166) for XBB.1.5 and 7.7 (n=4), 18.0 (n=37), 5.8 (n=13) and 27.6% (n=115) for BA.2.86 in SN2, SN3, SP2 and SH respectively. While no neutralization was observed against XBB.1.5 and BA.2.86, a low fraction of nAbs retained ADCP and ADCD activities in the SN2 (1.9 – 7.8%) and SP2 (2.2 – 6.5%) cohorts (**Fig. 1e, g**). Differently, a bigger fraction of nAbs in SN3 and SH retained both neutralization and Fc functions. Indeed, 15.0 – 35.4% of nAbs in the SN3 cohort were able to induce ADCP and ADCD against XBB.1.5 and BA.2.86, while 17.7 – 30.5% of antibodies retained these activities in the SH cohort (**Fig. 1f, h**). Next, we evaluated the Fc function potencies induced by nAbs in the four different cohorts and the S protein domains mainly involved in these activities (**Supplementary Fig. 3b-e**). No major differences were observed in the SN2 cohort for S protein trimer, RBD and NTD binding nAbs (**Supplementary Fig. 3b**). However, the low number of antibodies active against XBB.1.5 and BA.2.86 made this evaluation difficult. Differently, a third booster dose (SN3) or hybrid immunity (SP2) improved nAb Fc activities, despite losing potency against tested variants. In the SN3 cohort, NTD-binding nAbs showed the strongest ADCD activity while similar levels of activity are observed for ADCP (**Supplementary Fig. 3c**). In SP2, RBD- and NTD-binding nAbs showed similar potency which is higher than what observed for S protein trimer-targeting antibodies (**Supplementary Fig. 3d**). As for the SH cohort, nAbs binding the NTD showed the strongest ADCP and ADCD activities (**Supplementary Fig. 3e**).

### Antibody potency and breadth to alpha and beta human coronaviruses

To assess the breadth of neutralization to other alpha and beta human coronaviruses (h-CoV), in addition to SARS-CoV-1 and 2, we tested all nAbs isolated in the SH cohort against HKU-1, 229E and OC43. Initially, we evaluated the binding activity of all nAbs to the h-CoV S proteins. Our data showed that over 94.0% of nAbs were specific for SARS-CoV-2 only. Only, 3.8 (n=20), 1.0 (n=7), 0.3 (n=2) and 0.1% (n=1) of SARS-CoV-2 nAbs showed binding to SARS-CoV-1, OC43, 229E and HKU-1 respectively (**Supplementary Fig. 4a; Supplementary Table 4**). We next evaluated the neutralization activity and potency of cross-binding nAbs to SARS-CoV-1, OC43, 229E and HKU-1 pseudoviruses. Our results revealed that 45.0% (n=9) of SARS-CoV-1 S protein binding nAbs were neutralizing, with a 50% neutralization dose (ND_50_) ranging from 54.2 to 22,000.4 ng ml^-1^. Similarly, 57.1% (n=4) of nAbs reacting to OC43 S protein also showed neutralization activity but with an overall low potency with an ND_50_ ranging from 804.0 to 38,490.0 ng ml^-1^. The single nAb reacting to HKU-1 poorly neutralized the pseudovirus (ND_50_ 29,030.0 ng ml^-1^) while the two 299E S protein binding nAbs did not show neutralization activity against this alpha coronavirus (**Supplementary Fig. 4b**).

### Epitope mapping of cross-neutralizing nAbs

To understand the S protein regions mainly involved in cross-protection against the Omicron variants, we investigated the neutralization activity of RBD- and NTD-targeting nAbs (**Fig. 2**). RBD-targeting nAbs were classified based on their ability to compete with the Class 1/2 antibody J08^26^, the Class 3 antibody S309^27^, and the Class 4 antibody CR3022^28^, or for their lack of competition with the three tested antibodies (Not-competing)^13,25^. The neutralization activity against SARS-CoV-2 Wuhan virus in all cohorts was revealed to be mainly directed to the RBD Class 1/2 epitope region (**Fig. 2a-d**). In the SN2 group (*n* = 46), we observed a lack of neutralization activity against most Omicron variants tested except for one NTD-targeting nAb which neutralized both BF.7 and BQ.1.1 (**Fig. 2a**). In the SN3 cohort (n=197), after a third booster dose, we observed a more cross-reactive antibody response with 3.0 – 16.8% of nAbs able to neutralize all Omicron variants tested. The majority of these nAbs were directed against the RBD Class 1/2 epitope region (**Fig. 2b**). The SP2 cohort (n=215) showed 3.7 – 14.0% of cross-protection against Omicron BA.5, BA.2.75, BF.7 and BQ.1.1, while no neutralization was observed to XBB.1.5, EG.5.1.1 and BA.2.86. In this cohort, cross-neutralization was mainly driven by NTD and RBD Class 3 targeting nAbs (**Fig. 2c**). After a subsequent infection and additional vaccine dose, the antibody response in the SH cohort (n=413) showed a different profile. Cross-neutralization was observed against all tested Omicron variants with a frequency ranging from 6.5 to 62.7%. Of note, the majority of cross-neutralizing nAbs in the SH cohort were directed towards the RBD Class 1/2 region, changing the target profile compared to the less mature SP2 immune response (**Fig. 2d**). When we evaluated the neutralization potency, expressed as IC_100_, of cross-neutralizing nAbs, we observed in all cohorts that RBD binding antibodies were more potent than the NTD-targeting group (**Supplementary Fig. 5a-g; Supplementary Table 5**). As for the different RBD classes, our data showed that Class 3 nAbs were overall the most potent against Omicron variants in the SP2 cohort, showing up to 2.6- and 8.7-fold lower GM-IC_100_ compared to Class 1/2 and 4 nAbs respectively. In contrast, the most potent nAbs in the SN3 and SH cohorts were directed against the Class 1/2 epitope region. Overall, Class 1/2 nAbs in SN3 were up to 28.9-fold more potent than Class 3 antibodies, while in SH Class 1/2 nAbs had a GM-IC_100_ 4.7- and 13.0-fold lower compared to Class 3 and 4 antibodies respectively (**Supplementary Fig. 5a-g; Supplementary Table 5**). We also evaluated the ADCP and ADCD of all RBD classes and NTD binding nAbs against XBB.1.5 and BA.2.86. A very small fraction of nAbs in the SN2 cohort, despite not presenting neutralization activity, retained ADCD function (**Fig. 2e, left panel**). Nevertheless, the low number of nAbs makes it difficult to properly evaluate the Fc response in this cohort. Class 1/2 and NTD-binding nAbs were the most active in the SN3 cohort (**Fig. 2e, middle left panel**). Differently, Class 4 nAbs were the most active in the SP2 cohort, and Fc functions were further matured in SH individuals, where up to 85.7 and 57.1% of antibodies belonging to this class showed ADCP and ADCD function respectively (**Fig. 2e, middle right and right panels**). Further maturation of Class 1/2 and 3 in the SH cohort rescued the antibody Fc functions, expanding the fraction of nAbs able to induce ADCP and ADCD compared to SP2 (**Fig. 2e, middle right and right panels**). Our data also highlight the impact of the mutations present on the XBB.1.5 and BA.2.86 S proteins on antibody neutralization and Fc functions. Indeed, the unique set of mutations on the BA.2.86 S protein positioned in the Class 3 epitope region (N450D and K356T) seem to be extremely evasive in the SN3 cohort, leading to higher reduction of neutralization, ADCP and ADCD compared to XBB.1.5. Finally, nAbs in the SH cohort showed similar levels of neutralization and Fc functions against both XBB.1.5 and BA.2.86 (**Fig. 2e-g**).

**Fig. 2.**
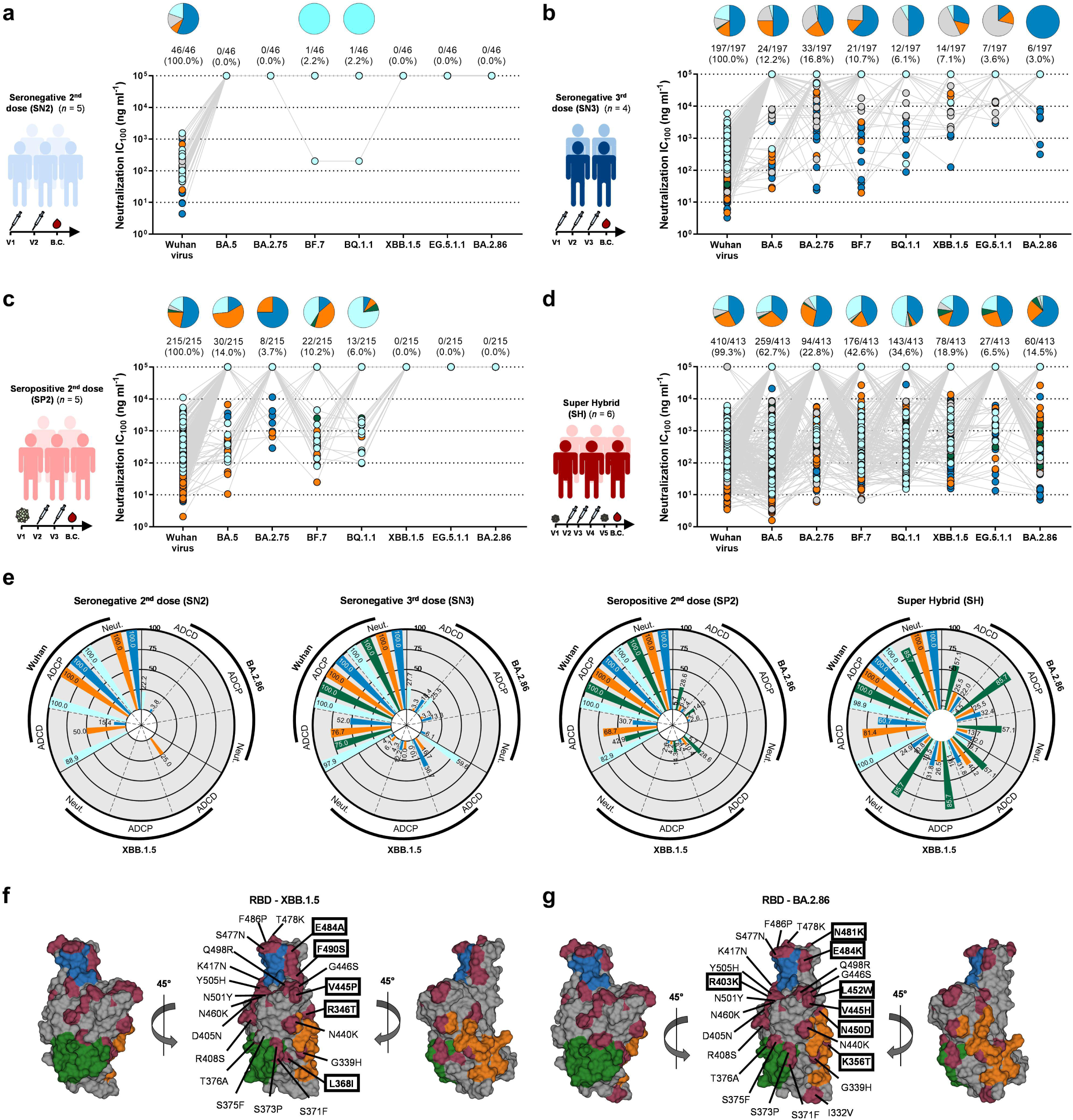
Distribution of RBD and NTD-targeting nAbs against Omicron variants. **a-d,** Pie charts show the distribution of cross-protective nAbs based on their ability to target Class 1/2 (blue), Class 3 (orange) and Class 4 (dark green) regions on the RBD, as well as not-competing nAbs (gray) and NTD-targeting nAbs (cyan). Dot charts show the neutralization potency, reported as IC_100_ (ng ml^-1^), of nAbs against the Wuhan virus and the Omicron BA.5, BA.2.75, BF.7, BQ.1.1, XBB.1.5, EG.5.1.1 and BA.2.86 variants observed in the SN2 (**a**), SN3 (**b**), SP2 (**c**) and SH (**d**) cohorts. The number and percentage of nAbs are denoted on each graph. **e,** Radar plots show the frequency of nAbs retaining neutralization, ADCP and ADCD activities against SARS-CoV-2 Wuhan, XBB.1.5 and BA.2.86. **f-g**, Representation of the SARS-CoV-2 RBD of XBB.1.5 (**f**) and BA.2.86 (**g**). In dark red are highlighted the mutations present on the RBD for both variants. Boxed mutated residue labels indicate unique mutations for the specific variant. Highlighted in blue, orange and green indicate Class 1/2, 3 and 4 epitope regions.

### B cell maturation and expansion in super hybrid immunity

In addition to the neutralization, Fc functions and epitope mapping analyses, we investigated the B cell expansion and maturation from hybrid (SP2 n sequences=278^7^) to super hybrid immunity (SH n sequences=441). Sequences were clustered by binning the clones to their inferred germlines (centroids) and according to 80% nucleotide sequence identity in the heavy complementary determining region 3 (CDRH3). Clusters were defined as antibody families including at least five or more members as previously described^25,29^. We found five immunoglobulin heavy variable (IGHV) and joining (IGHJ) rearrangements, IGHV1-24;IGHJ6-1, IGHV1-58;IGHJ3-1, IGHV3-53;IGHJ6-1, IGHV3-66;IGHJ4-1 and IGHV3-66;IGHJ6-1, to be predominantly expanded after an additional infection and vaccination dose. In addition, we observed six germlines, IGHV1-18;IGHJ4-1, IGHV1-69;IGHJ4-1, IGHV1-69;IGHJ5-1, IGHV2-70;IGHJ6-1, IGHV3-48;IGHJ4-1, IGHV5-10-1;IGHJ4-1, that emerged and expanded only in SH individuals broadening the B cell repertoire in this cohort (**Fig. 3a**). The five germlines expanded in SH, showed 1.20- to 2.67-fold higher somatic hypermutation (SHM) levels compared to SP2, with the IGHV1-58;IGHJ3-1 and IGHV3-53;IGHJ6-1 being the most mutated with a median V gene mutation of almost 9.0% (**Fig. 3b**). As for the six germlines found only in SH, we found B cells using the IGHV1-18;IGHJ4-1 rearrangement to be the most mutated with a median V gene mutations of 9.7%, while the IGHV5-10-1;IGHJ4-1 germline was the least mutated with a median mutation frequency of 3.9% (**Fig. 3c**). Next, we analyzed the binding and neutralization profiles of germlines expanded from SP2 or found only SH. The expanded germlines were found to mainly target the S protein RBD Class 1/2 epitope region with the exception of IGHV1-24;IGHJ6-1 which were found to preferentially bind the NTD (**Fig. 3d**). The neutralization data revealed that IGHV3-53;IGHJ6-1, IGHV3-66;IGHJ4-1 and IGHV3-66;IGHJ6-1 were the most cross-protective germlines against the SARS-CoV-2 variants tested, with IGHV3-53;IGHJ6-1 showing the highest potency with a GM-IC_100_ of 136.8 ng ml^-1^ (**Fig. 3e, left panel; Supplementary Table 6**). The germlines expanded only in SH also showed to be mainly directed against the S protein RBD but targeted different regions of the RBD. Indeed, IGHV1-69;IGHJ4-1, IGHV1-69;IGHJ5-1 targeted mainly the Class 1/2, while the germlines IGHV1-18;IGHJ4-1, IGHV2-70;IGHJ6-1, and IGHV5-10-1;IGHJ4-1 were directed almost exclusively towards the Class 3 epitope region. Interestingly, the germline IGHV3-48;IGHJ4-1 preferentially recognized the NTD and was shown to be the most cross-protective, covering 7 out 8 (87.5%) SARS-CoV-2 variants tested in this study with a GM-IC_100_ of 167.1 ng ml^-1^ (**Fig. 3d**). Conversely, the broadly expanded RBD-targeting germline IGHV1-69;IGHJ4-1 showed the lowest level of cross-protection despite some nAbs in this group being highly mutated and carrying almost 13% of mutations in the V gene (**Fig. 3e, right panel; Supplementary Table 6**).

**Fig. 3.**
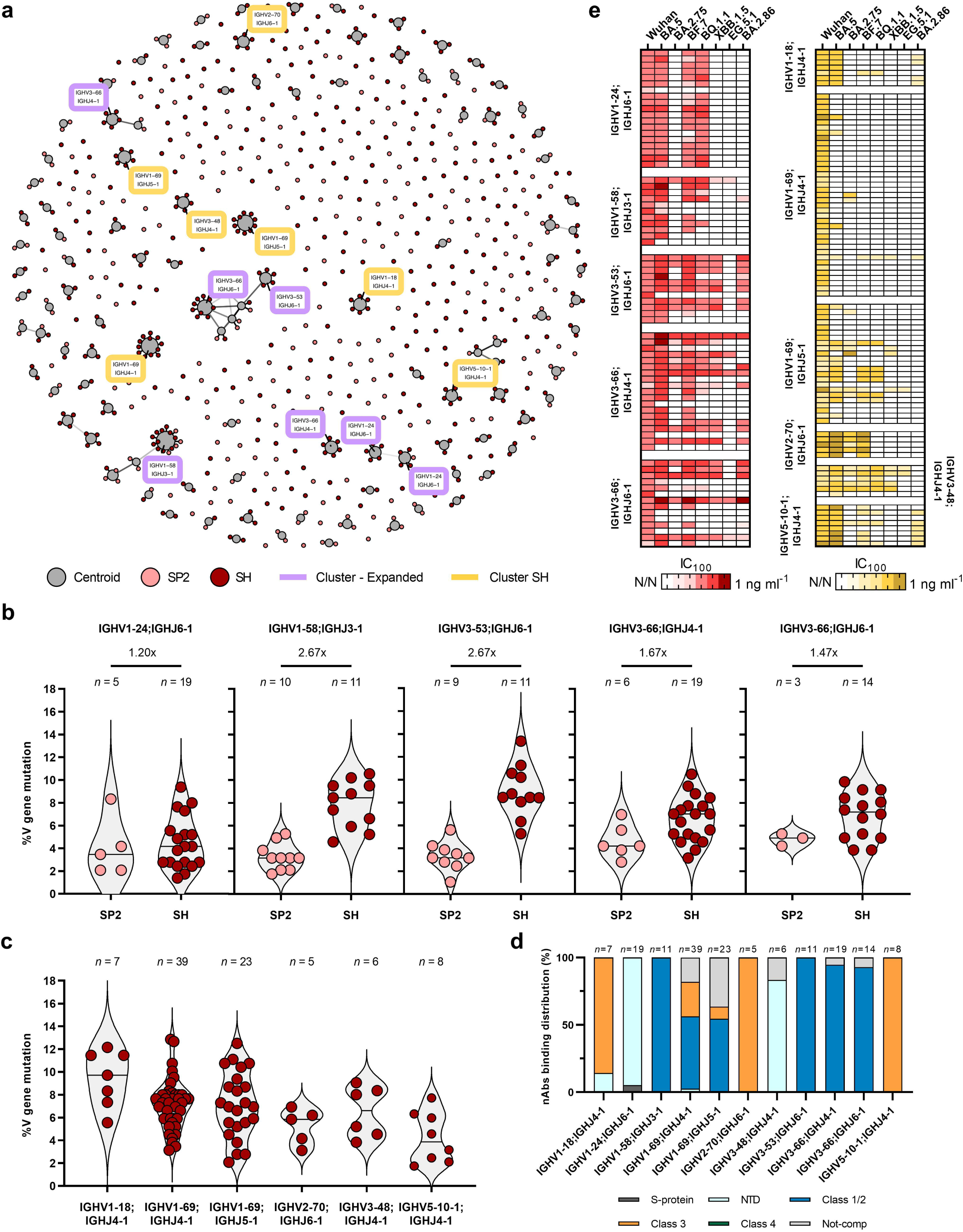
SH germlines expansion and characterization. **a,** Network plot shows the clonally-expanded antibody families in SP2 and SH. Centroids and nAbs from SP2 and SH groups are shown in gray, pink and dark red, respectively. Clusters and expanded clones are highlighted in gold and dark red, respectively. **b-c,** The violin plots show the V gene somatic mutation frequency of germlines expanded from SP2 to SH (**b**) or found exclusively in SH (**c**). The number of nAbs for each germline and fold-change are denoted on each graph. Violin plots show the median of V gene mutations. **d,** Bar graphs show the distribution of nAbs binding the S protein trimer (light gray), NTD (cyan) and RBD (dark gray) for expanded germlines. The number of nAbs per each germline is denoted on the graph. **e,** The heatmap shows the IC_100_ of predominant germlines expanded from SP2 to SH (left panel) or found exclusively in SH (right panel).

### B cell convergence after homologous or heterologous immunization

Next, we compared the evolution of the B cell repertoire after homologous or heterologous immunization in the SN3 (n sequences=289^25^) and SH cohorts. Interestingly, we observed a strong convergence of the B cell response in these two groups that shared the expansion of five different B cell rearrangements, IGHV1-58;IGHJ3-1, IGHV1-69;IGHJ3-1, IGHV1-69;IGHJ4-1, IGHV3-53;IGHJ6-1 and IGHV3-66;IGHJ6-1 (**Fig. 4a**). The five germlines, known to encode for potently neutralizing RBD-targeting Class 1 and Class 2 nAbs^13,28,30,31^, constituted 27.0 (n=78) and 22.7% (n=100) of the whole functional B cell repertoire in SN3 and SH respectively (**Supplementary Table 7**). While these germlines were predominantly expanded in both SN3 and SH, they showed different neutralization profiles against SARS-CoV-2 Wuhan and Omicron variants. Indeed, in the SN3 cohort, nAbs derived from IGHV1-69;IGHJ4-1 germlines were the most cross-reactive even if they showed a low to medium neutralization potency with a GM-IC_100_ of 2,065.1 ng ml^-1^ (**Fig. 4b, top panel; Supplementary Table 6**). Differently, in the SH cohort, IGHV1-69;IGHJ4-1 derived-nAbs were the least cross-reactive while antibodies encoded by the other four predominant germlines showed improved neutralization potency and breadth. The IGHV3-53;IGHJ6-1 and IGHV3-66;IGHJ6-1 were found to encode for the most cross-neutralizing nAbs with a GM-IC_100_ of 136.8 and 176.4 ng ml^-1^ respectively, which is 2.99- and 2.84-fold more potent of nAbs encoded by the same germlines in the SN3 cohort (**Fig. 4b, bottom panel; Supplementary Table 6**). These germlines showed broad neutralization against all variants including the highly mutated BA.2.86 (14/25; 56.0%), while only 2 out of 25 (8.0%) of these nAbs were able to cover the EG.5.1.1 variant which was predominant in Europe from July 2023 to November 2023, before the surge of JN.1. We then evaluated the Fc-functions induced by the five predominant germlines against the SARS-CoV-2 Wuhan, XBB.1.5 and BA.2.86 viruses and again observed different functional profiles. Indeed, IGHV1-69;IGHJ4-1 gene-derived nAbs in SN3 showed the strongest ADCP and ADCD compared to antibodies encoded by the same germlines in the SH cohort (**Fig. 4c, left panel**). IGHV3-53;IGHJ6-1 and IGHV3-66;IGHJ6-1 nAbs in SH showed the strongest Fc-functions in this cohort (**Fig. 4c, right panel; Supplementary Table 8**). When we analyzed the somatic hypermutation (SHM) levels of nAbs encoded by the five predominant germlines, we observed that only IGHV1-69;IGHJ4-1 nAbs had similar levels of V gene mutations between SN3 and SH. The remaining predominant germlines showed 1.24- to 1.57-fold higher levels of V gene mutations in SH with IGHV1-69;IGHJ3-1 and IGHV3-53;IGHJ6-1 being the most mutated (**Supplementary Fig. 6a**). In addition to shared germlines, we observed that SH had unique germlines not expanded in the SN3 cohort. The five most abundant were the IGHV1-18;IGHJ4-1, IGHV1-2;IGHJ4-1, IGHV1-24;IGHJ4-1, IGHV3-21;IGHJ6-1 and IGHV5-10-1;IGHJ4-1 (**Fig. 4a**). Interestingly, IGHV1-18;IGHJ4-1 and IGHV5-10-1;IGHJ4-1 were neither found in the SP2 cohort and were exclusive for the SH group. Antibodies from the IGHV1-24;IGHJ4-1, IGHV3-21;IGHJ6-1 and IGHV5-10-1;IGHJ4-1 germlines showed the highest cross-neutralization activity, covering 75% of tested SARS-CoV-2 variants with a GM-IC_100_ of 289.0, 287.1 and 91.3 respectively (**Supplementary Fig. 6b; Supplementary Table 7**). These three germlines were also the least mutated, with an average V gene mutations frequency below 6.0% suggesting space for further maturation of these nAbs (**Supplementary Fig. 6c**).

**Fig. 4.**
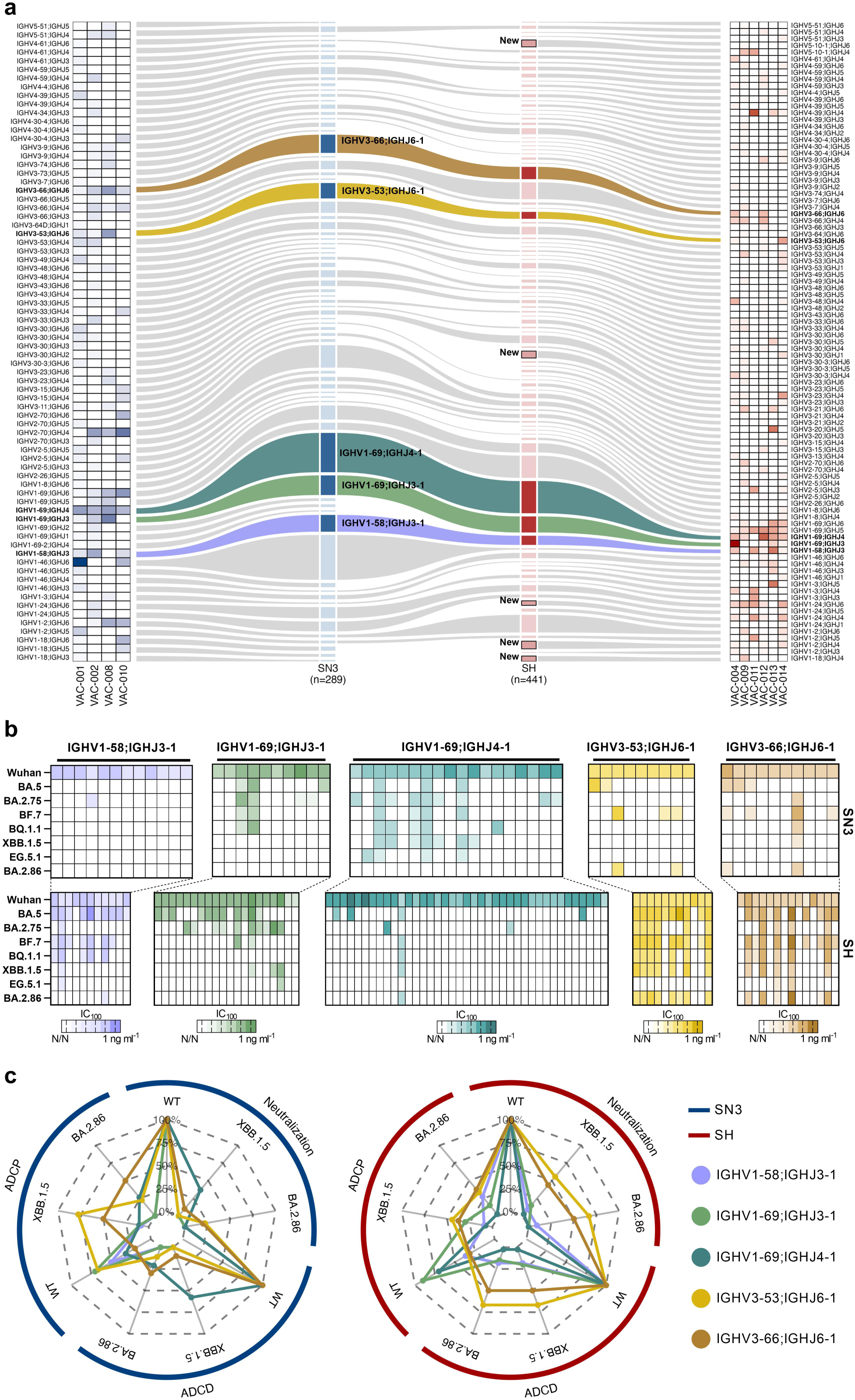
B cell repertoire and functional characterization of predominant germlines. **a,** Heatmaps and alluvial plots display the antibody IGHV;IGHJ gene rearrangements frequency for each single donor and for pulled nAbs respectively for SN3 (left panel) and SH (right panel). In the alluvial plots, the top five shared V-J gene rearrangements shared between SN3 and SH were highlighted. Selected germlines were highlighted as light purple, green, dark green, gold and brown for IGHV1-58;IGHJ3-1, IGHV1-69;IGHJ3-1, IGHV1-69;IGHJ4-1, IGHV3-53;IGHJ6-1 and IGHV3-66;IGHJ6-1 respectively. Box with black borders in the stratum identify the top five germlines expanded exclusively in SH. **b,** Heatmaps show the neutralization activity for the five most shared germlines between SN3 and SH against BA.5, BA.2.75, BF.7, BQ.1.1, XBB.1.5, EG.5.1.1 and BA.2.86. **c,** Radar plots describe the neutralization, ADCP, and ADCD activities of predominant germlines shared between SN3 and SH against Wuhan, XBB.1.5 and BA.2.86. The percentages of functionality for neutralization, ADCP, and ADCD are reported within each radar plot.

## DISCUSSION

In this work, we characterized in depth the antibody response of individuals with super hybrid immunity to understand how the B cell compartment matured in response to SARS-CoV-2 variants over time. We longitudinally analyzed a unique panel of almost 1,000 nAbs isolated from four different cohorts dissecting both homologous (SN2 and SN3) and heterologous (SP2 and SH) immune responses. Our data revealed that further maturation of the B cell compartment in SH led to the strongest and most cross-reactive antibody response to SARS-CoV-2 variants including the highly mutated BA.2.86. Interestingly, the antibody response in SH remained mainly focused on SARS-CoV-2 as less than 10% of nAbs showed cross-reactivity to other alpha and beta H-CoV. The most abundant class of nAbs after homologous immunization, independently from two or three mRNA vaccine doses, is the Class 1/2. Further maturation of the SP2 antibody response redirects nAbs from Class 3 and NTD to Class 1/2, converging with the antibody response observed after homologous immunization. Surprisingly, the main B cell germlines that stood behind the high levels of cross-protection observed in SH were antibodies encoded by the IGHV3-53/3-66 genes, which were highly evaded by all Omicron variants in less matured immunological responses like SN2, SN3 and SP2^13,14^. This observation suggests that the B cell compartment, after additional maturation, prefers to resiliently restore and expand germlines induced by the initial SARS-CoV-2 Wuhan S protein (i.e. the antigenic sin) over naïve B cells. Therefore, independently from the priming that individuals received after vaccination or infection, we observed a strong convergence in the antibody response. Of note, up to 56% of IGHV3-53/3-66 encoded nAbs were able to neutralize the highly mutated BA.2.86 variant, while only 8% of these nAbs cross-protected against EG.5.1.1. This observation highlights the importance of the highly conserved residue Y33 placed in the heavy chain complementary determining region 1 (H-CDR1) of IGHV3-53/3-66 encoded nAbs as it was previously shown to form extensive hydrophobic interactions with the RBD residue F456 mutated exclusively on the EG.5.1.1 lineage (F456L)^17,30^. In addition, this observation explains, at single cell level, why the highly mutated BA.2.86 did not become predominant worldwide despite exhibiting substantial antigenic drift, remarkably enhanced receptor binding affinity, fusogenicity and infectivity to lung cells compared to other variants^32–34^. Anyway, BA.2.86 rapidly evolved and at the end of 2023 the JN.1 sublineage emerged becoming the predominant variant worldwide. JN.1 carries only one additional mutation in the RBD, L455S, compared to the ancestral BA.2.86 variant which resulted in decreased ACE2 affinity but enhanced immune evasion^35,36^. Future work will help to understand the classes of antibodies evaded and the germlines that are still able to retain neutralization against JN.1. Indeed, we identified low mutated B cell germlines uniquely expanded in SH (IGHV1-18;IGHJ4-1 and IGHV5-10-1;IGHJ4-1), suggesting a stretch of the antibody repertoire which could be rapidly deployed against future SARS-CoV-2 variants including JN.1. Overall, our work provides unique information on the longitudinal evolution of the B cell compartment in response to SARS-CoV-2 variants highlighting similarities and differences between homologous and heterologous vaccination, and how the imprinting of the antigenic sin restored and drove antibody maturation.

## METHODS

### Enrollment of donors with super hybrid immunity and human sample collection

Human samples from individuals with SH, of both sexes, were collected through a collaboration with the Azienda Ospedaliera Universitaria Senese, Siena (IT). All subjects enrolled gave their written consent. The study that allowed the enrollment of subjects in all three cohorts was approved by the Comitato Etico di Area Vasta Sud Est (CEAVSE) ethics committees (Parere 17065 in Siena, amendment 13 December 2021) and conducted according to good clinical practice in accordance with the declaration of Helsinki (European Council 2001, US Code of Federal Regulations, ICH 1997). This study was unblinded and not randomized. Six subjects were enrolled in this cohort. Subjects in the SH cohort were exposed two times to SARS-CoV-2 infection and received three or four mRNA vaccine doses. First infections occurred between October 2020 and July 2022, while second infections occurred between January and December 2022. Vaccinations occurred between December 2020 and December 2022. Given the exploratory nature of this study, no statistical methods were used to predetermine sample size.

### Single cell sorting of SARS-CoV-1 and SARS-CoV-2 S-protein^+^ memory B cells from COVID-19 vaccinees

Peripheral blood mononuclear cells (PBMCs) isolation was performed as previously described^6,7,25^. Briefly, PBMCs were isolated from heparin-treated whole blood by density gradient centrifugation (Ficoll-Paque PREMIUM, SigmaAldrich). After separation, cells were incubated at room temperature for 20 minutes with the viability dye Live/Dead Fixable Aqua (Invitrogen; Thermo Scientific). Then, cells were washed with PBS and incubated with 50 µL of 20% normal rabbit serum (Life technologies) diluted in PBS to saturate unspecific bindings. After 30 minutes of incubation at 4°C, cells were washed with PBS and stained with SARS-CoV-1 S-protein labeled with Strep-Tactin™XT DY-649 (iba-lifesciences cat# 2-1568-050) and SARS-CoV-2 S-protein labeled with Strep-TactinXT DY-488 (iba-lifesciences cat# 2-1562-050) for 30 min at 4°C. Then, a surface staining was performed using CD19 V421 (BD cat# 562440), IgM PerCP-Cy5.5 (BD cat# 561285), CD27 PE (BD cat# 340425), IgD-A700 (BD cat# 561302), CD3 PE-Cy7 (BioLegend cat# 300420), CD14 PE-Cy7 (BioLegend cat# 301814), CD56 PECy7 (BioLegend cat# 318318). After 30 minutes of incubation at 4°C, stained memory B cells were single cell-sorted with a BD FACSAria™ Fusion (BD Biosciences) into 384-well plates and were incubated for 14 days with IL-2, IL-21 and irradiated 3T3-CD40L as previously described^37^.

### SARS-CoV-2 authentic viruses neutralization assay

All SARS-CoV-2 authentic virus neutralization assays were performed in the biosafety level 3 (BSL3) laboratories at Toscana Life Sciences in Siena (Italy), which is approved by a Certified Biosafety Professional and inspected annually by local authorities. To assess the neutralization potency and breadth of nAbs against live SARS-CoV-2 and its variants, a cytopathic effect-based microneutralization assay (CPE-MN) was performed as previously described^6,7,13,25^. Briefly, 100 median Tissue Culture Infectious Dose (100 TCID_50_) of SARS-CoV-2 virus was co-incubated with nAbs for 1 hour at 37°C, 5% CO_2_. The virus-antibody mixtures were then moved into a 96-well plate containing a sub-confluent Vero E6 cell monolayer. Plates were incubated for 3-4 days at 37°C in a humidified environment with 5% CO_2_, then examined for CPE by means of an inverted optical microscope by two independent operators. Single cell sorting supernatants were tested at single point dilution to identify positive hits and antibody to be recombinantly expressed through transcriptionally active polymerase chain reaction (TAP-PCR). TAP supernatant, used to evaluate the neutralization potency of identified nAbs, were tested at a starting dilution of 1:4 and diluted step 1:2. Single replicate and technical duplicates were performed to evaluate single cell sorting supernatant and to evaluate the IC_100_ of TAP respectively. In each plate positive and negative control were used as previously described^6,7,13,25^.

### SARS-CoV-2 virus variants CPE-MN neutralization assay

The SARS-CoV-2 viruses used to perform the CPE-MN neutralization assay were the Wuhan (SARS-CoV-2/INMI1-Isolate/2020/Italy: MT066156), Omicron BA.5 (GISAID ID: EPI_ISL_13389618), BA.2.75 (GISAID ID: EPI_ISL_14732896), BF.7 (GISAID ID: EPI_ISL_13499917), BQ.1.1 (GISAID ID: EPI_ISL_15455664), XBB.1.5 (GISAID ID: EPI_ISL_17272995), EG.5.1.1 (GISAID ID: EPI_ISL_18245523) and BA.2.86 (GISAID ID: EPI_ISL_18221650). The BA.2.86 strain (hCoV-19/France/IDF-IPP17625/2023) was supplied by the National Reference Centre for Respiratory Viruses hosted by Institut Pasteur (Paris, France). The human sample from which strain hCoV-19/France/IDF-IPP17625/2023 was isolated has been provided by Dr Aude LESENNE from Cerballiance Ile De France Sud, Lisses.

### Single cell RT-PCR and Ig gene amplification and transcriptionally active PCR expression

To express our nAbs as full-length IgG1, 5 µL of cell lysate from the original 384-cell sorting plate were used for reverse transcription polymerase chain reaction (RT-PCR), and two rounds of PCRs (PCRI and PCRII-nested) as previously described^6,7,25^. Obtained PCRII products were used to recover the antibody heavy and light chain sequences, through Sanger sequencing, and for antibody cloning into expression vectors as previously described^6,7,25^. TAP-PCR reaction was performed using 5 μL of Q5 polymerase (NEB), 5 μL of GC Enhancer (NEB), 5 μL of 5X buffer,10 mM dNTPs, 0.125 µL of forward/reverse primers and 3 μL of ligation product, using the following cycles: 98°/2’, 35 cycles 98°/10’’, 61°/20’’, 72°/1’ and 72°/5’. TAP products were purified under the same PCRII conditions, quantified by Qubit Fluorometric Quantitation assay (Invitrogen) and used for transient transfection in Expi293F cell line following manufacturing instructions.

### Expression and purification XBB.1.5, BA.2.86 and H-CoV S proteins

The plasmids encoding SARS-CoV-2 6P WT and for the seven S proteins of the season H-CoV was generously provided by Prof. Jason S. McLellan. All proteins were expressed and purified as previously described^6^. Briefly, plasmids encoding for XBB.1.5, BA.2.86 and H-CoV S proteins were transiently transfected in ExpiCHO-S cells (Thermo Fisher) using ExpiFectamine™ CHO Reagent. Cells were grown for six days at 37°C with 8% CO_2_ in shaking conditions at 125 rpm according to the manufacturer’s protocol (Thermo Fisher). ExpiFectamine™ CHO Enhancer and ExpiCHO™ Feed were added 18 to 22 hours post-transfection to boost transfection, cell viability, and protein expression. Both types of cell cultures (for the spike of SARS-CoV-2 and H-CoVs) were harvested five days after transfection and the proteins were purified by immobilized metal affinity chromatography (FF Crude) followed by dialysis into final buffer. Cell culture supernatants were clarified by centrifugation (1,200x g, 30 min, 4°C) followed by filtration through a 0.45 μm filter. Chromatography purification was conducted at room temperature using ÄKTA Go purifier system from GE Healthcare Life Sciences. Specifically, filtered culture supernatant was purified with a 5 mL HisTrap FF Crude column (GE Healthcare Life Sciences) previously equilibrated in Buffer A (20 mM NaH2PO4, 500 mM NaCl + 30 mM Imidazole pH 7.4). The flow rate for all steps of the HisTrap FF Crude column purification was 5 ml/min. The culture supernatant of each spike protein was applied to a single 5 mL HisTrap FF Crude column. The column was washed in Buffer A with 4 column volumes (CV). spike proteins were eluted from the column by applying a first step elution of 5CV of 60% Buffer B (20 mM NaH_2_PO_4_, 500 mM NaCl + 500 mM Imidazole pH 7.4). Elution fractions were collected in 1 ml each and analyzed by SDS-PAGE. Fractions containing the S protein were pooled and dialyzed against PBS buffer pH 7.4 using Slide-A-Lyzer™ Dialysis Cassette 10K MWCO (Thermo Scientific) overnight at 4°C. The dialysis buffer used was at least 200 times the volume of the sample. The final spike protein concentration was determined by measuring absorbance at 562 nm using Pierce™ BCA Protein Assay Kit (Thermo Scientific™). Proteins were dispensed into 0.5 ml aliquots and stored at -80°C.

### ELISA assay with SARS-CoV-2 NTD, RBD and S2 subunits

To determine the binding specificity of nAbs to the SARS-CoV-2 S protein domains we performed and ELISA to the RBD, NTD and S2 domains. The assay was performed as previously described^7,25^. Briefly, 3 µg ml^-1^ of SARS-CoV-2 subunits diluted in carbonate-bicarbonate buffer (E107, Bethyl Laboratories), were coated in 384-well plates (microplate clear, Greiner Bio-one), and blocked with 50µl/well of blocking buffer (phosphate-buffered saline, 1% BSA) for 1h at 37 °C. After washing (phosphate-buffered saline and 0.05% Tween-20), plates were incubated with mAbs diluted 1:5 in dilution buffer (phosphate-buffered saline, 1% BSA, 0.05% Tween-20) and step-diluted 1:2 in dilution buffer. Anti-Human IgG-Peroxidase antibody (Fab specific) produced in goat (Sigma) diluted 1:45,000 in dilution buffer for RBD and NTD plates, while 1:80,000 in dilution buffer for S2 plates, was added and incubated for 1h at 37 °C. Plates were then washed, incubated whit TMB substrate (Sigma) for 15 min before adding 25 µl/well of stopping solution (H_2_SO4 0.2M). The OD values were identified using the Varioskan Lux Reader (Thermo Fisher Scientific) at 450 nm. Each condition was tested in duplicate and samples were considered positive if the OD value was two-fold the blank.

### ELISA assay with H-CoVs S proteins

Antibody binding specificity against the H-CoVs SARS-CoV-1, 229E, OC43 and HKU-1 S proteins was detected by ELISA as previously described^13^. Briefly, 384-well plates (microplate clear, Greiner Bio-one) were coated with 3 µg ml^-1^ of streptavidin (Thermo Fisher) diluted in carbonate-bicarbonate buffer (E107, Bethyl Laboratories) and incubated at RT overnight. The next day, plates were incubated for 1h at RT with 3 µg ml^-1^ of H-CoVs S proteins, and saturated with 50 µl/well of blocking buffer (phosphate-buffered saline, 1% BSA) for 1h at 37 °C. Following, 25 µl/well of mAbs diluted 1:5 in dilution buffer (phosphate-buffered saline, 1% BSA, 0.05% Tween-20) were added and serially diluted 1:2 and then incubated for 1h at 37 °C. Finally, 25 µl/well of alkaline phosphatase-conjugated goat Anti-Human IgG diluted 1:2,000 in dilution buffer were added. mAbs binding to the S proteins were detected using 25 µl/well of PNPP (p-nitrophenyl phosphate; Thermo Fisher) and the reaction was measured at a wavelength of 405nm using the Varioskan Lux Reader (Thermo Fisher Scientific). After each incubation step, plates were washed with washing buffer (phosphate-buffered saline and 0.05% Tween-20). Sample buffer was used as a blank and the threshold for sample positivity was set at two-fold the OD of the blank. Technical duplicates were performed.

### SARS-CoV-1 pseudotype based microneutralization assay

To screen single cell sorting supernatants and identify mAbs able to neutralize SARS-CoV-1 we performed a microneutralization assay in 384 well-plates. The HEK293TN-hACE2 cell line was generated by lentiviral transduction of HEK293TN (System Bioscience, Cat#LV900A-1) cells as described in Notarbartolo S. et al^38^. SARS-CoV-1 lentiviral pseudotype particles were generated as described in Conforti et al. using the SARS-CoV1 SPIKE plasmid pcDNA3.3_CoV1_D28 (Addgene plasmid # 170447)^39^. HEK293TN-hACE2 were plated 10,000/well in 384-well flat plates (Corning cat#3765) using DMEM medium complete (10% FBS, 2mM L-glutamine, 1% Pen-strep,1mM Sodium pyruvate, 1% Non-Essential Amino Acid). After 24h, cells were infected with 0.1 MOI of SARS-CoV-1 pseudotyped viruses that were previously incubated with 7 µl of each mAb supernatant. Supernatants were diluted 1:5 with a pseudotyped viral solution at 0.1 multiplicity of infection (MOI) and were tested at single point dilution. PBS and the mAb S309 (tested starting from 10 µg ml^-1^ and serially diluted step 1:3) were used as negative and positive controls respectively. The mix was incubated for 1h at 37°C and then 25 µl were added into 50 µl of HEK293TN-hACE2 pre-seeded wells. Plates were incubated at 37°C for 24h after which luciferase activity was measured reading the plate on Varioskan™ LUX (Thermo Scientific) using the Bright-Glo Luciferase Assay System (Promega), according to the manufacturer’s recommendations. Percent of inhibition was calculated relative to pseudotype virus-only control. 50% neutralization dose (ND_50_) values were established by nonlinear regression using Prism v.8.1.0 (GraphPad).

### H-CoV (229E, HKU-1 and OC43) pseudotype based microneutralization assays

The neutralization assay for 229E, HKU-1 and OC43 H-CoV pseudotyped viruses was performed as described for SARS-CoV-1 pseudotype virus using 96-well plates. In addition, appropriate cell lines expressing H-CoVs receptor were used to allow infection and evaluation of neutralization activity of isolated nAbs. Specifically, HUH7, CHO-K1 and HEKTN/17 cell lines were used for 229E, HKU-1 and OC43 respectively. Briefly, 100 µl containing 1×10^4^ cells were plated in each well in white 96-well plates using DMEM medium complete (10% FBS, 2mM L-glutamine, 1% Pen-strep,1mM Sodium pyruvate, 1% Non-Essential Amino Acid). 24h later, cells were infected with 0.1 MOI of pseudotyped viruses that were previously co-incubated for 1h at 37°C with serial dilution of TAP supernatants in 50 µl. A 7-point dose response curve was obtained by diluting TAP supernatants step 1:3. After 24 incubation at 37°C for 24h, luciferase activity was measured reading the plate on Varioskan™ LUX (Thermo Scientific™) using the Bright-Glo Luciferase Assay System (Promega), according to the manufacturer’s recommendations. Percent inhibition was calculated relative to pseudotype virus-only control. ND_50_ (Neutralization Dose) values were established by nonlinear regression using Prism v.8.1.0 (GraphPad). The average ND_50_ value for each antibody was determined from a minimum of three independent technical replicates and two independent experiments. Technical triplicates were performed.

### Flow cytometry-based competition assay

To characterize mAb candidates based on their interaction with S protein epitopes, a flow cytometry-based competition assay was performed. As previously described^7^, 1 mg of magnetic beads (Dynabeads His-Tag, Invitrogen) were coupled with 200 µg of histidine-tagged SARS-CoV-2 S protein. Then, S protein-beads (20 μg ml^-1^) were incubated with unlabeled neutralizing antibodies at room temperature for 40 minutes and the sample was subsequently washed in PBS-1% BSA. To evaluate the S protein epitope competition, antibodies RBD Class 1/2 (J08), Class 3 (S309), Class 4 (CR3022) binders or NTD (4A8) binders were labelled with different fluorophores (Alexa Fluor 647, 488, 594) using the Alexa Fluor NHS Ester kit (Thermo Scientific). Next, fluorescent labeled antibodies were incubated with S protein-beads for 40 minutes at RT. After incubation, the S protein-antibodies mix was washed with PBS, resuspended in 150 μL of PBS-BSA 1%, and analyzed using BD FACSymphony™ A3 (BD Biosciences). As positive and negative controls, beads with or without S protein incubated with labeled antibodies were used. FACSDiva Software (version 9) and FlowJo (version 10) were used for data acquisition and analysis, respectively.

### Measurement of ADCP and ADCD functions triggered by neutralizing antibodies

Antibody-dependent cellular phagocytosis (ADCP) was performed using a Flow cytometry–based assay. As previously described^40^, stabilized histidine-tagged SARS-CoV-2 S protein (Wuhan, XBB.1.5, BA.2.86) was labelled with Strep-Tactin™XT Conjugate DY-649 (IBA Lifesciences) and conjugated to magnetic beads according to the manufacturer’s instructions. The mix S protein-beads was incubated with nAbs for 1 h at RT and then mixed with monocytic THP-1 cell line (50.000 per well). After 18h of incubation at 37°C, THP-1 cells were washed with PBS and fixed with fixation buffer (BioLegend) following the manufacturer’s guidelines. Next, cells resuspended in 100 μL of PBS1X were acquired by BD FACSymphony™ A3 (BD Biosciences). FlowJo software (version 10) was used for data analysis and phagocytosis was evaluated as percentage of fluorescent beads engulfed by THP-1 multiplied by the median fluorescence intensity of the population. To explore the antibody dependent complement deposition (ADCD), Expi293F cells (Thermo Fisher, Cat#A14527) were transiently transfected with SARS-CoV-2 original S protein, XBB.1.5, or BA.2.86 expression vectors (pcDNA3.1_ spike_del19) using the ExpiFectamine Enhancer (Thermo Fisher)^40^. After 48h, monoclonal antibodies were incubated with S protein-expressing cells at 37 °C, with 5% CO_2_ and 120 rpm shaker speed for 30 min. Then, 6% of baby rabbit complement (Cedarlane) diluted in Expi medium was added, and cells were incubated at 37 °C, with 5% CO_2_ and 120 rpm shaker speed. After 30 minutes of incubation, cells were incubated with goat anti-rabbit polyclonal antibody against C3-FITC conjugated (MP Biomedicals) for 1 h on ice. Then, stained cells were fixed with fixation buffer (BioLegend) for 15 min on ice and resuspended in 100 μL of PBS. BD FACSymphony™ A3 (BD Biosciences) was used for data acquisition and results were reported as median fluorescence intensity of the FITC signal detected.

### Functional repertoire analyses

nAbs VH and VL sequence reads were manually curated and retrieved using CLC sequence viewer (Qiagen). Aberrant sequences were removed from the data set. Analyzed reads were saved in FASTA format and the repertoire analyses was performed using Cloanalyst (http://www.bu.edu/computationalimmunology/research/software/)^41,42^.

### Network plot of clonally expanded antibody families

A network map was built by representing each clonal family with a centroid and connecting centroids sharing a similar sequence. The centroid sequence was computed with Cloanalyst to represent the average CDRH3 sequence for each clonal family, and Hamming distance was calculated for each antibody CDRH3 sequence to represent the relationship within the clonal family. Levenshtein distance was calculated between each centroid representative of each clonal family to investigate the relationship between clonal families. Levenshtein distance was calculated with the R package stringdistm v0.9.8 (https://cran.r-project.org/web/packages/stringdist/index.html) and normalized between 0 and 1. A network graph was generated with the R package ggraph v2.0.5 (https://ggraph.data-imaginist.com/index.html) with Fruchterman-Reingold layout algorithm and the figure was assembled with ggplot2 v3.3.5. The size of the centroid is proportional to the number of antibodies belonging to the same clonal family, while the color of each node represents the antibody origin: dark blue for seronegative 3rd dose, and dark red for superhybrids.

### Alluvial plot of germline frequency distribution

An alluvial plot was generated to display the frequency distribution of IGHV;IGHJ germlines among the two analyzed cohorts: seronegative 3rd dose (SN3) and superhybrids (SH). The cohorts are represented as two separate categories (strata), and the germline frequency for each single cohort is represented by the flow size. The analysis included all antibodies with fully sequenced VH chains, totaling 289 entries for SN3 and 441 entries for SH. The change in IGHV;IGHJ germline frequency amoung the two cohorts was evaluated, and the five germlines with higher, unaltered frequency were highlighted. Specifically, germline IGHV3−66;IGHJ6, IGHV3−53;IGHJ6, IGHV1−69;IGHJ4, IGHV1−69;IGHJ3, and IGHV1−58;IGHJ3 were selected. Additionally, germlines not observed in the SN3 cohort but comprising more than five entries in the SH cohort (IGHV5−10−1;IGHJ4, IGHV3−21;IGHJ6, IGHV1−2;IGHJ4, IGHV1−24;IGHJ4, IGHV1−18;IGHJ4) were highlighted within the SH stratum using a black rectangle. Spider plots were created for each cohort to depict the functionality of the selected germlines in terms of neutralization, antibody-dependent cellular phagocytosis (ADCP), and antibody-dependent complement deposition (ADCD). The percentage of functional antibodies associated with each germline was determined using predefined thresholds of 100,000, 100,000, and 4,000 for neutralization, ADCP, and ADCD, respectively. The figure was assembled with ggplot2 v3.3.5. All the scripts and the data is available via github (https://github.com/dasch-lab/SARS-CoV-2_superhybrid).

### Statistical analysis

Statistical analysis was assessed with GraphPad Prism Version 8.0.2 (GraphPad Software, Inc., San Diego, CA). Nonparametric Mann-Whitney t test was used to evaluate statistical significance between the two groups analyzed in this study. Statistical significance was shown as * for values ≤ 0.05, ** for values ≤ 0.01, and *** for values ≤ 0.001.

## Acknowledgments

This work received funding by the European Research Council (ERC) advanced grant (agreement number 101098201 (PROACTIVE)). P.M. acknowledges support from the Research Foundation Flanders (COVID19 research grant G0H4420N) and Internal Funds KU Leuven (grant 3M170314). Work in O.S. laboratory is funded by Institut Pasteur, Urgence COVID-19 Fundraising Campaign of Institut Pasteur, Fondation pour la Recherche Médicale (FRM), ANRS, the Vaccine Research Institute (ANR-10-LABX-77), Labex IBEID (ANR-10-LABX-62-IBEID), ANR / FRM Flash Covid PROTEO-SARS-CoV-2, ANR Coronamito, HERA european funding, Sanofi and IDISCOVR. The E.S.-L. laboratory is funded by Institut Pasteur, the INCEPTION program (Investissements d’Avenir grant ANR-16-CONV-0005), the Labex IBEID (grant no. ANR-10-LABX-62-IBEID), the HERA Project DURABLE (grant no 101102733) and the NIH PICREID (grant no U01AI151758). N.T. and M.M.N. acknowledge support from Wellcome Trust 360G-Wellcome-220981_Z_20_Z.

## Author contributions

Conceived the study: R.R. and E.A.; Provided PBMCs and enrolled SH donors: S.P., M.F., I.R., M.T. and F.M.; Isolated PBMCs and performed single cell sorting: I.P.; Provided SARS-CoV-2 viruses: P.M. E;S;-L. and O.S.; Expanded and titrated SARS-CoV-2 variants: G.P.; Performed neutralization assays in BSL3 facilities: I.P., G.P., E.P. and G.A.; SARS-CoV-1 and H-CoV pseudotype production and neutralization assay: E.P., M.M.N.; Expression of recombinant S proteins: E.P. and G.A.; Performed antibody expression: I.P., G.A., G.R. and F.P.; Performed ELISA assay on SARS-CoV-1, SARS-CoV-2 and H-CoV S protein, and SARS-CoV-2 domains: G.P. and G.A.; Performed epitope mapping: I.P.; Performed ADCP and ADCD assays: I.P.; Recovered VH and VL sequences and performed the repertoire analyses: P.P., G.M. D.C. and E.A.; Manuscript writing: R.R. and E.A.; Final revision of the manuscript: I.P., G.P., E.P., G.A., P.P., G.M., D.C., G.R., F.P., M.M.N., S.P., M.F., I.R., M.T., F.M., D.M., P.M., N.T., O.S., R.R. and E.A.; Coordinated the project: E.A..

## Additional information

**Correspondence and requests for materials** should be addressed to E.A.

## Data availability

Source data are provided with this paper. All data supporting the findings in this study are available within the article or can be obtained from the corresponding author upon request.

## SUPPLEMENTARY FIGURES

**Supplementary Fig. 1.**
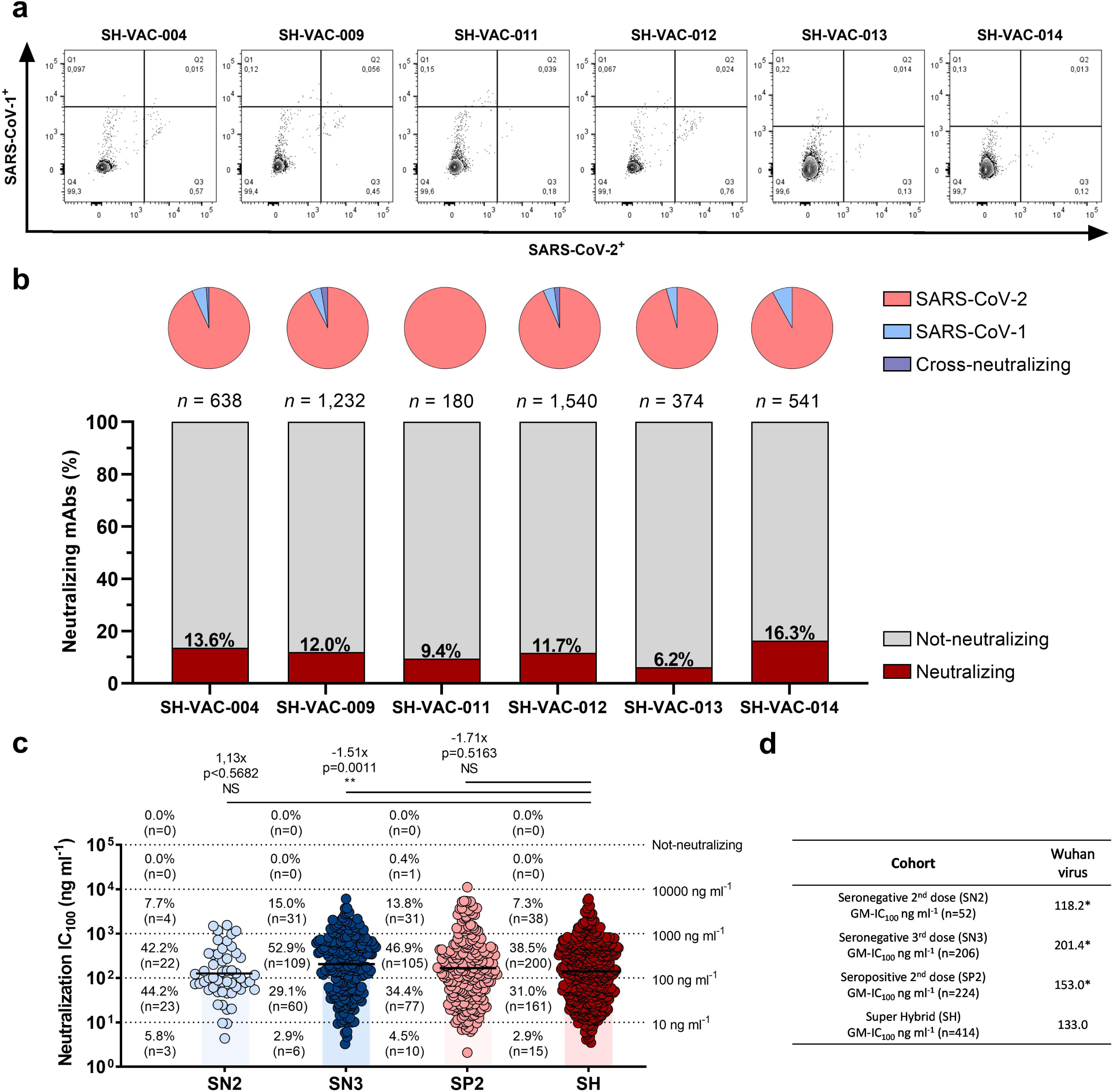
B cell frequencies and identification of SARS-CoV nAbs. **a,** Gate strategy used to single cell sort antigen specific CD19^+^CD27^+^IgD^-^IgM^-^ class-switched memory B cells for all donors in the SH cohort. **b,** The bar chart at the bottom of the graph shows the percentage of nAbs identified per each donor in the SH cohort. The neutralizing and not-neutralizing antibody fractions are represented in dark red and light gray respectively. The number of antibodies for each donor is denoted on the graph. Pie charts at the top of the graph represent the distribution of SARS-CoV-1 (light blue), SARS-CoV-2 (pink) and cross-neutralizing (violet) antibodies in all SH donors. **c,** Scatter dot charts show the neutralization potency, reported as IC_100_ (ng ml^-1^), of nAbs tested against the original Wuhan SARS-CoV-2 virus for SN2, SN3, SP2 and SH shown in light blue, blue, pink, and dark red respectively. The number, percentage, GM-IC_100_ (black lines and colored bars), fold-change and statistical significance of nAbs are denoted on each graph. Reported fold-change and statistical significance are in comparison with the SH cohort. Technical duplicates were performed for each experiment. A nonparametric Mann–Whitney t test was used to evaluate statistical significances between groups. Two-tailed p-value significances are shown as *p < 0.05, **p < 0.01, and ***p < 0.001. **d,** Table summarizing the GM-IC_100_ values against the Wuhan virus for all tested cohorts. Asterisked values were reported in previous publications^7,25^.

**Supplementary Fig. 2.**
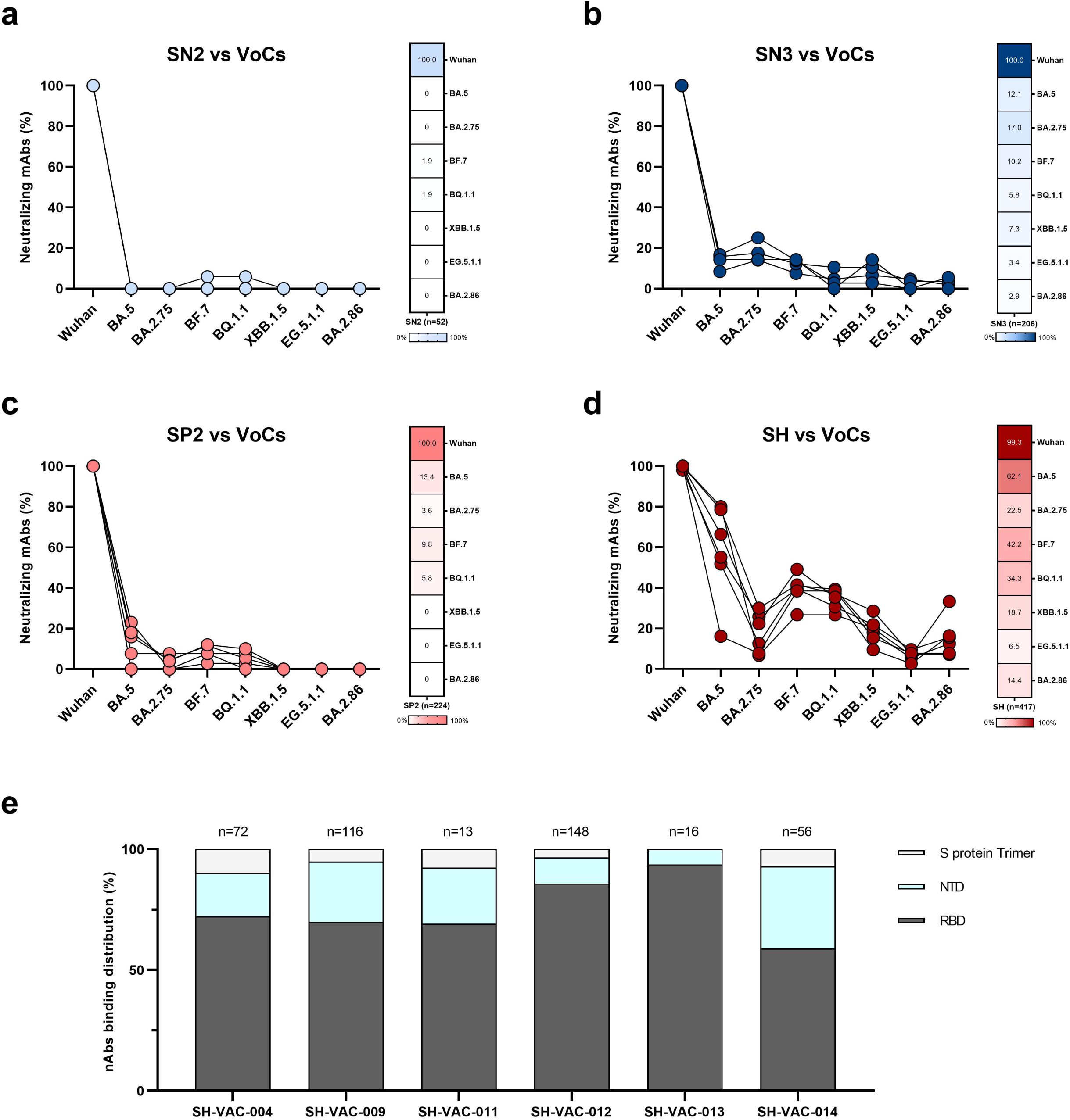
nAbs cross-neutralization and binding distribution. **a-d,** Graphs show the fold change percentage of nAbs in SN2 (**a**), SN3 (**b**), SP2 (**c**) and SH (**d**) against BA.5, BA.2.75, BF.7, BQ.1.1, XBB.1.5, EG.5.1.1 and BA.2.86 Omicron variants compared to the original Wuhan SARS-CoV-2 virus. The heatmaps show the overall percentage of nAbs able to neutralize tested SARS-CoV-2 variants. **e,** The bar graph shows the percentage of S protein trimer (light gray), NTD (cyan) and RBD (dark gray) binding nAbs for each individual in the SH cohort. The number (n) of nAbs tested per each cohort is denoted on the graph.

**Supplementary Fig. 3.**
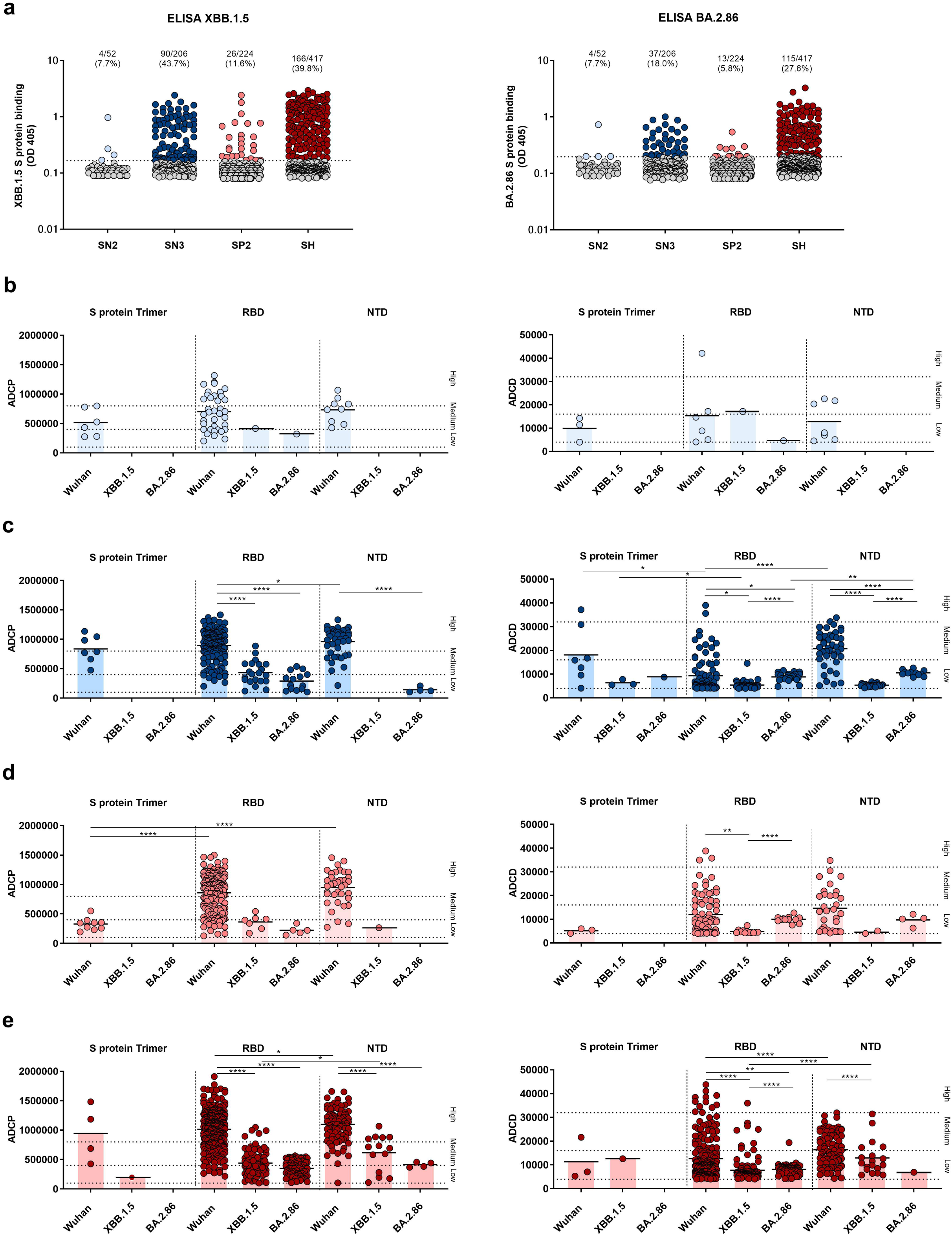
Binding and Fc functions to XBB.1.5 and BA.2.86. **a,** Scatter dot plots show the binding of nAbs isolated in all cohorts for XBB.1.5 (left panel) and BA.2.86 (right panel) S proteins. **b-e**, Graphs show the ADCP and ADCD potency between RBD, NTD and the S protein in trimeric conformation against SARS-CoV-2 original Wuhan virus, the Omicron XBB.1.5 and BA.2.86 variants in SN2 (**b**), SN3 (**c**), SP2 (**d**) and SH (**e**) cohorts. Non-parametric Mann–Whitney t-test was used to evaluate the statistical significance between groups. Two-tailed p value significances are shown as *p<0.05, **p < 0.01, ***p < 0.001, and ****p<0.0001.

**Supplementary Fig. 4.**
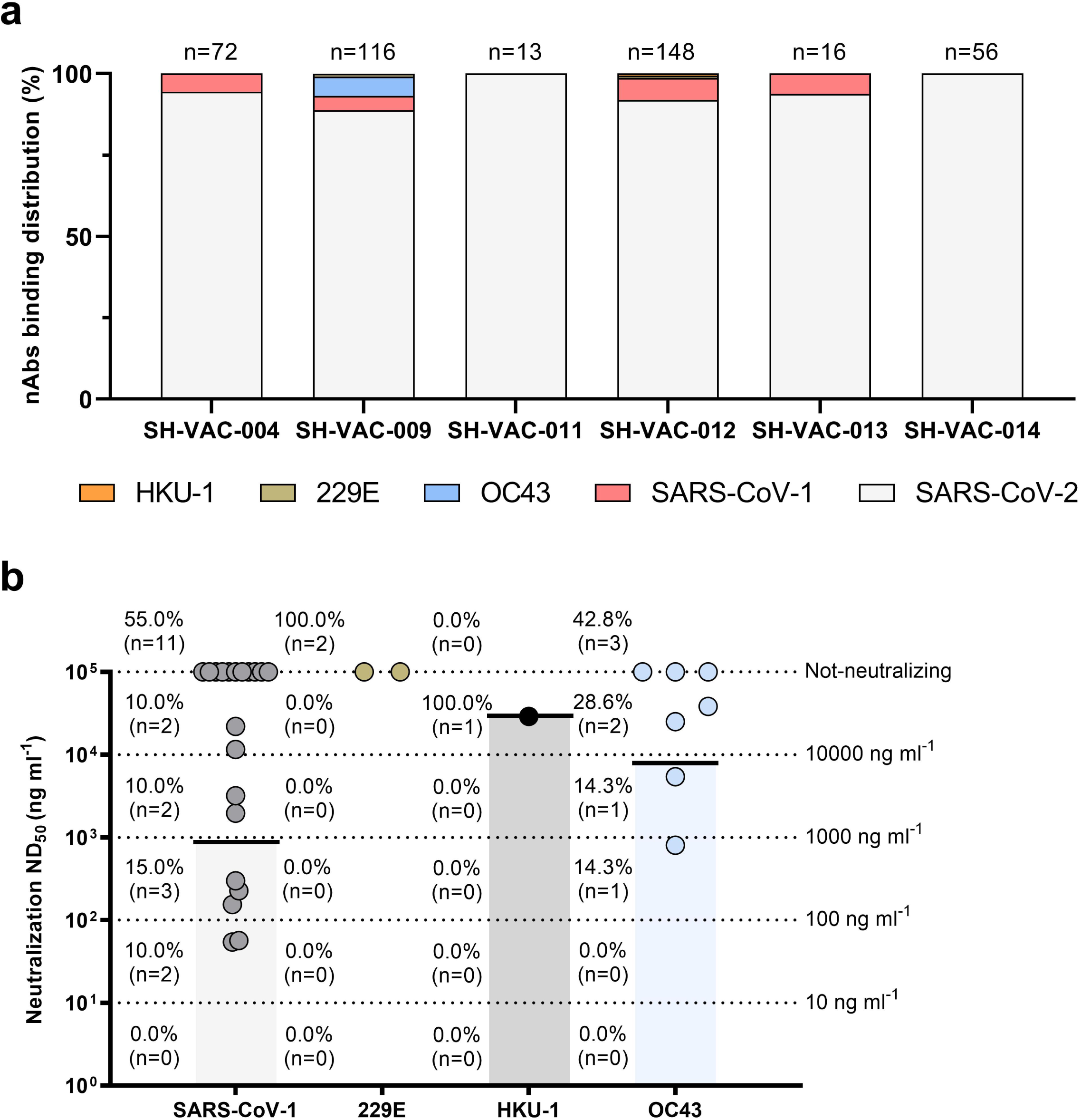
Binding and neutralization activity to alpha and beta H-CoV. **a,** The bar graph shows the percentage of nAbs binding to SARS-CoV-2 (light gray), SARS-CoV-1 (pink), OC43 (light blue), 229E (light brown) and HKU-1 (orange) for all donors tested in the SH cohort. **b,** Scatter dot charts show the neutralization potency, reported as IC_100_ (ng ml^-1^), of nAbs tested against SARS-CoV-1, 229E, HKU-1 and OC43. The number, percentage and GM-IC_100_ (black lines and colored bars) are denoted on each graph. Technical duplicates were performed for each experiment.

**Supplementary Fig. 5.**
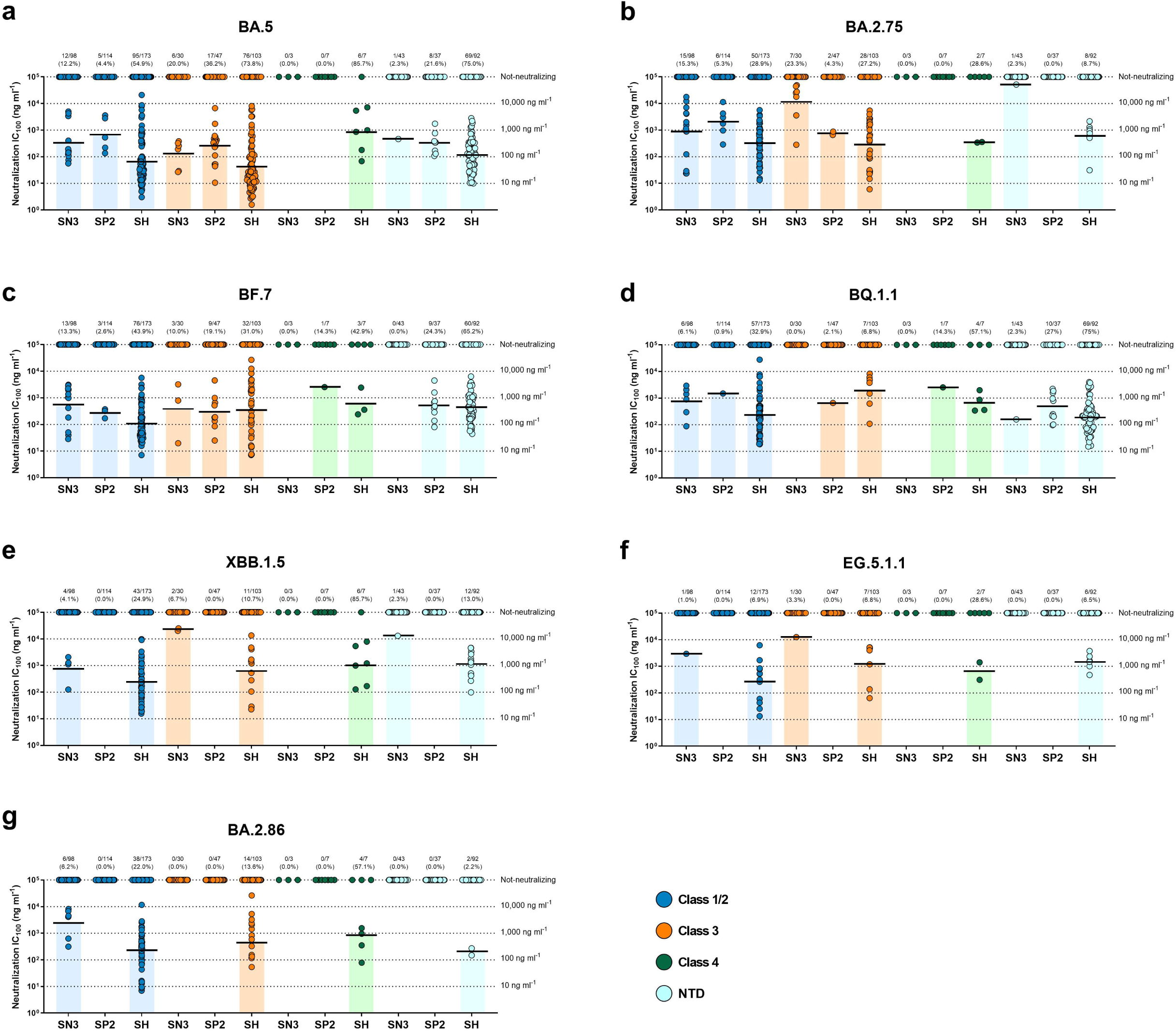
Neutralization potency of RBD Class 1/2, 3 and 4, and NTD-targeting nAbs. **a-g,** Scatter dot charts show the neutralization potency, reported as GM-IC_100_ (ng ml^-1^), of nAbs tested against SARS-CoV-2 Omicron variants BA.5 (**a**), BA.2.75 (**b**), BF.7 (**c**), BQ.1.1 (**d**), XBB.1.5 (**e**), EG.5.1.1 (**f**) and BA.2.86 (**g**). The number, percentage and GM-IC_100_ (black lines and colored bars) are denoted on each graph. Antibodies targeting the RBD Class 1/2, 3 and 4, and NTD are shown in blue, orange, green and cyan respectively.

**Supplementary Fig. 6.**
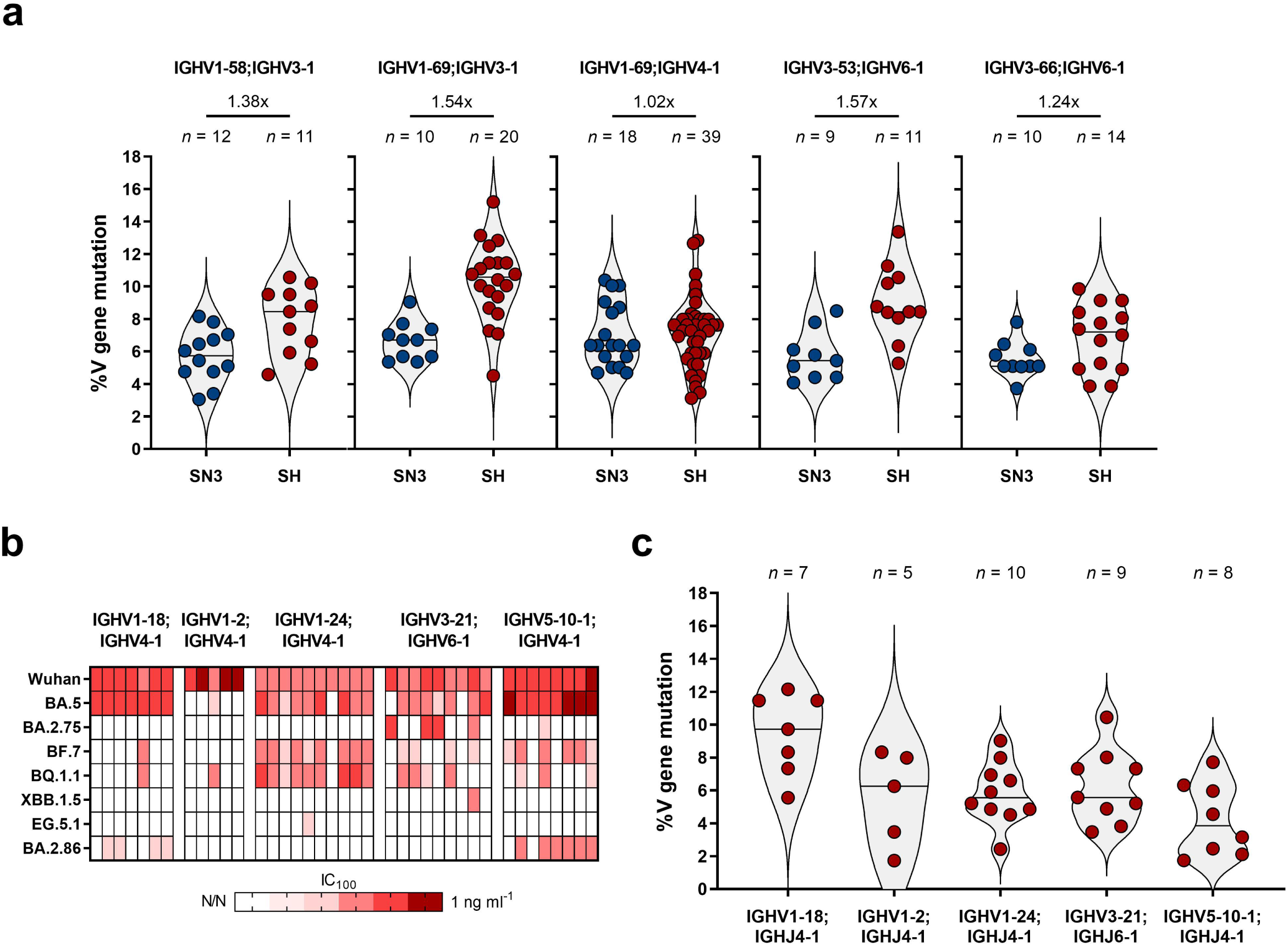
Characterization of SN3-SH shared germlines and SH exclusive germlines. **a,** The violin plots show the V gene somatic mutation frequency of IGHV1-24;IGHJ6-1, IGHV1-58;IGHJ3-1, IGHV3-53;IGHJ6-1, IGHV3-66;IGHJ4-1 and IGHV3-66;IGHJ6-1 gene derived nAbs. The number of nAbs for each germline and fold-change are denoted on each graph. Violin plots show the median of V gene mutations. **b,** The heatmap shows the IC_100_ of germlines found in SH but not in SN3 against all SARS-CoV-2 variants tested in this study. **c,** The violin plot shows the V gene somatic mutation frequency of selected germlines found exclusively in the SH cohort. The number of nAbs for each germline is denoted on each graph. Violin plots show the mean of V gene mutations.

## SUPPLEMENTARY TABLES

**Supplementary Table 1.**
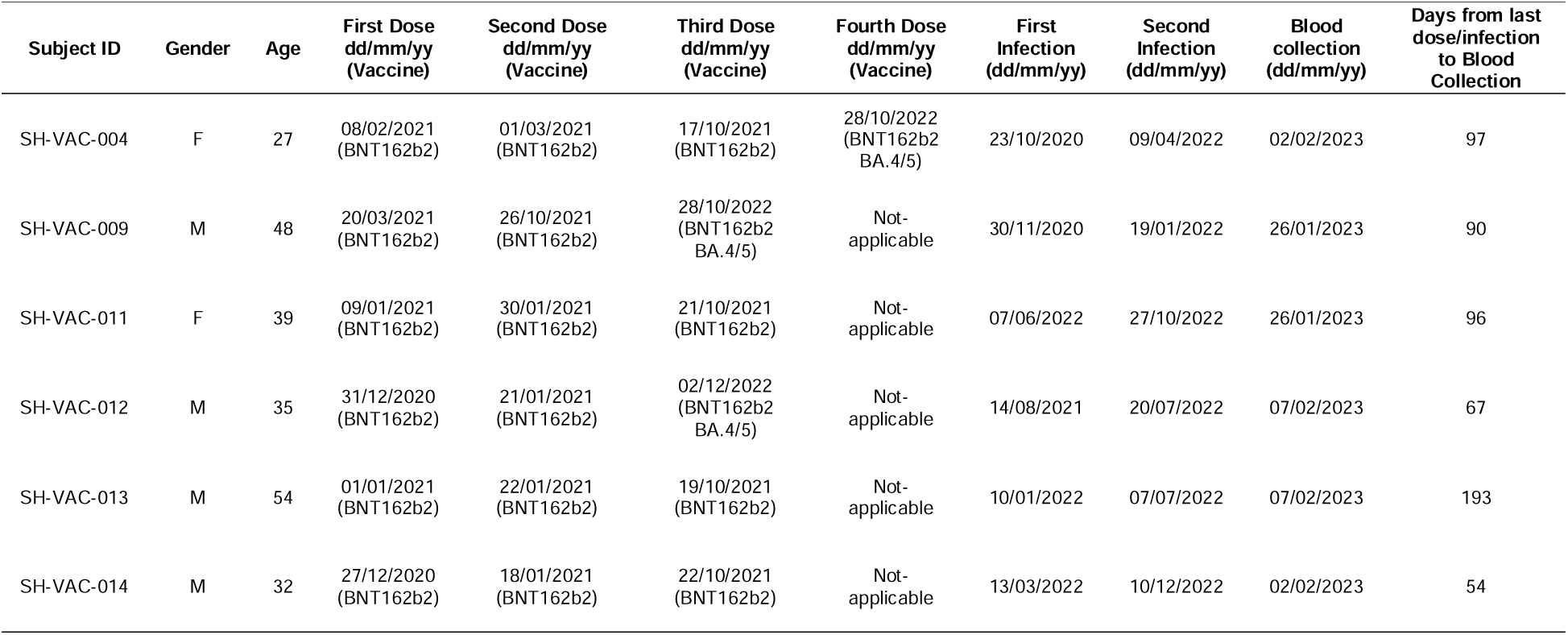
Clinical details of super hybrid immunity donors.

**Supplementary Table 2.** Frequencies, numbers and neutralization of antigen specific memory B cells in donors with super hybrid immunity.

| Subject ID | Frequency MBCs – SARS-CoV-1 spike (%) | Frequency MBCs – SARS-CoV-2 spike (%) | Frequency MBCs – Double positive (%) | Total Number MBCs Sorted | Total Number Neutralizing mAbs | Neutralizing SARS-CoV-2 (%) | Neutralizing SARS-CoV-1 (%) | Cross-neutralizing (%) |
| --- | --- | --- | --- | --- | --- | --- | --- | --- |
| VAC-004 | 0.097 | 0.57 | 0.015 | 638 | 87 | 81 (93.1) | 4 (4.6) | 2 (2.3) |
| VAC-009 | 0.12 | 0.45 | 0.056 | 1,232 | 148 | 137 (92.6) | 7 (4.7) | 4 (2.7) |
| VAC-011 | 0.15 | 0.18 | 0.039 | 180 | 17 | 17 (100.0) | 0 (0.0) | 0 (0.0) |
| VAC-012 | 0.067 | 0.76 | 0.024 | 1,540 | 182 | 169 (92.9) | 8 (4.4) | 5 (2.7) |
| VAC-013 | 0.22 | 0.13 | 0.014 | 374 | 23 | 22 (95.7) | 1 (4.3) | 0 (0.0) |
| VAC-014 | 0.13 | 0.12 | 0.013 | 541 | 88 | 81 (92.0) | 7 (8.0) | 0 (0.0) |

**Supplementary Table 3.** Neutralization potency (GM-IC_100_) in SN2, SN3, SP2 and SH.

| Cohort | Wuhan virus | Omicron BA.5 | Omicron BA.2.75 | Omicron BF.7 | Omicron BQ.1.1 | Omicron XBB.1.5 | Omicron EG.5.1.1 | Omicron BA.2.86 |
| --- | --- | --- | --- | --- | --- | --- | --- | --- |
| Seronegative 2<sup>nd</sup> dose (SN2)<br>GM-IC<sub>100</sub> ng mL<sup>-1</sup> (n=52) | 118.2 | N/A | N/A | 202.1 | 202.1 | N/A | N/A | N/A |
| Seronegative 3<sup>rd</sup> dose (SN3)<br>GM-IC<sub>100</sub> ng mL<sup>-1</sup> (n=206) | 201.4 | 525.0 | 3,712.4 | 1,046.6 | 1,664.1 | 5,474.4 | 7,994.1 | 2,427.9 |
| Seropositive 2<sup>nd</sup> dose (SP2)<br>GM-IC<sub>100</sub> ng mL<sup>-1</sup> (n=224) | 153.0 | 328.1 | 1,589.1 | 399.1 | 599.7 | N/A | N/A | N/A |
| Super Hybrid (SH)<br>GM-IC<sub>100</sub> ng mL<sup>-1</sup> (n=417) | 133.0 | 77.0 | 318.2 | 222.9 | 241.0 | 437.3 | 625.4 | 307.0 |

**Supplementary Table 4.** Binding specificity to SARS-CoV-2 S protein domains and other human betacoronaviruses.

| Subject ID | Total Number expressed nAbs | S protein binding nAbs (%) | RBD binding nAbs (%) | NTD binding nAbs (%) | Binding SARS-CoV-1 (%) | Binding OC43 (%) | Binding 229E (%) | Binding HKU-1 (%) |
| --- | --- | --- | --- | --- | --- | --- | --- | --- |
| VAC-004 | 72 | 7 (9.7) | 52 (72.2) | 13 (18.1) | 4 (5.6) | 0 (0.0) | 0 (0.0) | 0 (0.0) |
| VAC-009 | 116 | 6 (5.2) | 81 (69.8) | 29 (25.0) | 5 (4.3) | 7 (6.0) | 1 (0.9) | 0 (0.0) |
| VAC-011 | 13 | 1 (7.7) | 9 (69.2) | 3 (23.1) | 0 (0.0) | 0 (0.0) | 0 (0.0) | 0 (0.0) |
| VAC-012 | 148 | 5 (3.4) | 127 (85.8) | 16 (10.8) | 10 (6.8) | 0 (0.0) | 1 (0.7) | 1 (0.7) |
| VAC-013 | 16 | 0 (0.0) | 15 (93.7) | 1 (6.2) | 1 (6.2) | 0 (0.0) | 0 (0.0) | 0 (0.0) |
| VAC-014 | 56 | 4 (7.1) | 33 (58.9) | 19 (33.9) | 0 (0.0) | 0 (0.0) | 0 (0.0) | 0 (0.0) |

**Supplementary Table 5.** Neutralization potency of RBD Class 1/2, 3 and 4, and NTD nAbs.

| Cohort | Class 1/2 | Class 3 | Class 4 | NTD |
| --- | --- | --- | --- | --- |
| **BA.5** |  |  |  |  |
| Seronegative 3<sup>rd</sup> dose (SN3)<br>GM-IC<sub>100</sub> ng ml<sup>-1</sup> | 317.7 | 129.5 | N/A | 464.2 |
| Seropositive 2<sup>nd</sup> dose (SP2)<br>GM-IC<sub>100</sub> ng ml<sup>-1</sup> | 692.4 | 262.5 | N/A | 330.3 |
| Super Hybrid (SH)<br>GM-IC<sub>100</sub> ng ml<sup>-1</sup> | 66.1 | 42.3 | 857.4 | 115.7 |
| **BA.2.75** |  |  |  |  |
| Seronegative 3<sup>rd</sup> dose (SN3)<br>GM-IC<sub>100</sub> ng ml<sup>-1</sup> | 878.1 | 11,534.8 | N/A | 51,453.7 |
| Seropositive 2<sup>nd</sup> dose (SP2)<br>GM-IC<sub>100</sub> ng ml<sup>-1</sup> | 2,030.8 | 761.4 | N/A | N/A |
| Super Hybrid (SH)<br>GM-IC<sub>100</sub> ng ml<sup>-1</sup> | 332.6 | 276.1 | 348.9 | 611.7 |
| **BF.7** |  |  |  |  |
| Seronegative 3<sup>rd</sup> dose (SN3)<br>GM-IC<sub>100</sub> ng ml<sup>-1</sup> | 555.9 | 367.4 | N/A | N/A |
| Seropositive 2<sup>nd</sup> dose (SP2)<br>GM-IC<sub>100</sub> ng ml<sup>-1</sup> | 272.8 | 291.5 | 2537.2 | 505.0 |
| Super Hybrid (SH)<br>GM-IC<sub>100</sub> ng ml<sup>-1</sup> | 107.3 | 347.0 | 593.2 | 453.1 |
| **BQ.1.1** |  |  |  |  |
| Seronegative 3<sup>rd</sup> dose (SN3)<br>GM-IC<sub>100</sub> ng ml<sup>-1</sup> | 762.0 | N/A | N/A | 158.0 |
| Seropositive 2<sup>nd</sup> dose (SP2)<br>GM-IC<sub>100</sub> ng ml<sup>-1</sup> | 1,487.5 | 657.7 | 2,537.2 | 469.7 |
| Super Hybrid (SH)<br>GM-IC<sub>100</sub> ng ml<sup>-1</sup> | 238.6 | 1,897.6 | 674.3 | 185.6 |
| **XBB.1.5** |  |  |  |  |
| Seronegative 3<sup>rd</sup> dose (SN3)<br>GM-IC<sub>100</sub> ng ml<sup>-1</sup> | 774.5 | 22,412.6 | N/A | 13,017.0 |
| Seropositive 2<sup>nd</sup> dose (SP2)<br>GM-IC<sub>100</sub> ng ml<sup>-1</sup> | N/A | N/A | N/A | N/A |
| Super Hybrid (SH)<br>GM-IC<sub>100</sub> ng ml<sup>-1</sup> | 240.9 | 642.4 | 1,015.1 | 1,122.0 |
| **EG.5.1.1** |  |  |  |  |
| Seronegative 3<sup>rd</sup> dose (SN3)<br>GM-IC<sub>100</sub> ng ml<sup>-1</sup> | 2,954.4 | 12,542.5 | N/A | N/A |
| Seropositive 2<sup>nd</sup> dose (SP2)<br>GM-IC<sub>100</sub> ng ml<sup>-1</sup> | N/A | N/A | N/A | N/A |
| Super Hybrid (SH)<br>GM-IC<sub>100</sub> ng ml<sup>-1</sup> | 268.1 | 1,262.6 | 662.3 | 1,470.6 |
| **BA.2.86** |  |  |  |  |
| Seronegative 3<sup>rd</sup> dose (SN3)<br>GM-IC<sub>100</sub> ng ml<sup>-1</sup> | 2,427.9 | N/A | N/A | N/A |
| Seropositive 2<sup>nd</sup> dose (SP2)<br>GM-IC<sub>100</sub> ng ml<sup>-1</sup> | N/A | N/A | N/A | N/A |
| Super Hybrid (SH)<br>GM-IC<sub>100</sub> ng ml<sup>-1</sup> | 238.6 | 715.3 | 449.3 | 200.5 |

**Supplementary Table 6.** GM-IC_100_ ng ml^-1^ of SN3 and SH germlines.

| **SN3** | **GM-IC<sub>100</sub> ng ml<sup>-1</sup>** |
| --- | --- |
| IGHV1-58;IGHJ3-1 | 111.6 |
| IGHV1-69;IGHJ3-1 | 198.0 |
| IGHV1-69;IGHJ4-1 | 2065.1 |
| IGHV3-53;IGHJ6-1 | 408.8 |
| IGHV3-66;IGHJ6-1 | 500.2 |
| **SH** | **GM-IC<sub>100</sub> ng ml<sup>-1</sup>** |
| IGHV1-18;IGHJ4-1 | 90.1 |
| IGHV1-2;IGHJ4-1 | 65.0 |
| IGHV1-24;IGHJ4-1 | 289.0 |
| IGHV1-58;IGHJ3-1 | 167.0 |
| IGHV1-69;IGHJ3-1 | 71.7 |
| IGHV1-69;IGHJ4-1 | 101.4 |
| IGHV1-69;IGHJ5-1 | 165.1 |
| IGHV2-70;IGHJ6-1 | 14.1 |
| IGHV3-21;IGHJ6-1 | 287.1 |
| IGHV3-48;IGHJ4-1 | 167.1 |
| IGHV3-53;IGHJ6-1 | 136.8 |
| IGHV3-66;IGHJ4-1 | 165.3 |
| IGHV3-66;IGHJ6-1 | 176.4 |
| IGHV5-10-1;IGHJ4-1 | 91.3 |

**Supplementary Table 7.**
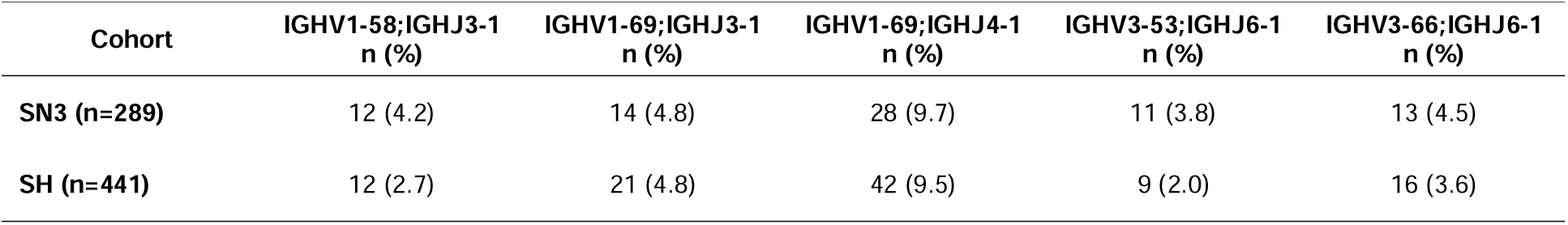
Abundance expanded germlines in SN3 and SH.

| Cohort | IGHV1-58;IGHJ3-1<br>n (%) | IGHV1-69;IGHJ3-1<br>n (%) | IGHV1-69;IGHJ4-1<br>n (%) | IGHV3-53;IGHJ6-1<br>n (%) | IGHV3-66;IGHJ6-1<br>n (%) |
| --- | --- | --- | --- | --- | --- |
| **SN3 (n=289)** | 12 (4.2) | 14 (4.8) | 28 (9.7) | 11 (3.8) | 13 (4.5) |
| **SH (n=441)** | 12 (2.7) | 21 (4.8) | 42 (9.5) | 9 (2.0) | 16 (3.6) |

**Supplementary Table 8.** Neutralization and Fc functions of predominant germlines.

| Cohort | IGHV;IGHJ | nAbs (n) | Neut.<br>WT (%) | Neutr.<br>XBB.1.5<br>(%) | Neutr.<br>BA.2.86<br>(%) | ADCP<br>WT<br>(%) | ADCP<br>XBB.1.5<br>(%) | ADCP<br>BA.2.86<br>(%) | ADCD<br>WT<br>(%) | ADCD<br>XBB.1.5<br>(%) | ADCD<br>BA.2.86<br>(%) |
| --- | --- | --- | --- | --- | --- | --- | --- | --- | --- | --- | --- |
| SH | IGHV1-58;IGHJ3 | 11 | 100.0 | 9.1 | 9.1 | 100.0 | 18.2 | 18.2 | 36.4 | 9.1 | 27.3 |
| SH | IGHV1-69;IGHJ3 | 20 | 100.0 | 15.0 | 0.0 | 100.0 | 20.0 | 15.0 | 90.0 | 30.0 | 15.0 |
| SH | IGHV1-69;IGHJ4 | 40 | 100.0 | 2.5 | 0.0 | 100.0 | 2.5 | 2.5 | 70.0 | 7.5 | 2.5 |
| SH | IGHV3-53;IGHJ6 | 9 | 100.0 | 55.6 | 66.7 | 100.0 | 66.7 | 66.7 | 33.3 | 44.4 | 33.3 |
| SH | IGHV3-66;IGHJ6 | 16 | 100.0 | 43.8 | 50.0 | 100.0 | 50.0 | 50.0 | 31.3 | 37.5 | 37.5 |
| SN3 | IGHV1-58;IGHJ3 | 12 | 100.0 | 0.0 | 0.0 | 100.0 | 0.0 | 0.0 | 50.0 | 8.3 | 0.0 |
| SN3 | IGHV1-69;IGHJ3 | 10 | 100.0 | 0.0 | 0.0 | 100.0 | 0.0 | 0.0 | 70.0 | 10.0 | 0.0 |
| SN3 | IGHV1-69;IGHJ4 | 19 | 100.0 | 36.8 | 0.0 | 100.0 | 57.9 | 21.1 | 31.6 | 10.5 | 26.3 |
| SN3 | IGHV3-53;IGHJ6 | 9 | 100.0 | 0.0 | 22.2 | 100.0 | 0.0 | 11.1 | 66.7 | 77.8 | 22.2 |
| SN3 | IGHV3-66;IGHJ6 | 10 | 100.0 | 10.0 | 20.0 | 100.0 | 10.0 | 30.0 | 20.0 | 50.0 | 50.0 |

